# A cortical locus for modulation of arousal states

**DOI:** 10.1101/2024.05.24.595859

**Authors:** Nithik Chintalacheruvu, Anagha Kalelkar, Jöel Boutin, Vincent Breton-Provencher, Rafiq Huda

**Affiliations:** WM Keck Center for Collaborative Neuroscience, Department of Cell Biology and Neuroscience, Rutgers University – New Brunswick, Piscataway, New Jersey, USA; Department of Psychiatry and Neuroscience, CERVO Brain Research Center, Université Laval, Québec City, Québec, Canada

## Abstract

Fluctuations in global arousal are key determinants of spontaneous cortical activity and function. Several subcortical structures, including neuromodulatory nuclei like the locus coeruleus (LC), are involved in the regulation of arousal. However, much less is known about the role of cortical circuits that provide top-down inputs to arousal-related subcortical structures. Here, we investigated the role of a major subdivision of the prefrontal cortex, the anterior cingulate cortex (ACC), in arousal modulation. Pupil size, facial movements, heart rate, and locomotion were used as non-invasive measures of arousal and behavioral state. We designed a closed loop optogenetic system based on machine vision and found that real time inhibition of ACC activity during pupil dilations suppresses ongoing arousal events. In contrast, inhibiting activity in a control cortical region had no effect on arousal. Fiber photometry recordings showed that ACC activity scales with the magnitude of spontaneously occurring pupil dilations/face movements independently of locomotion. Moreover, optogenetic ACC activation increases arousal independently of locomotion. In addition to modulating global arousal, ACC responses to salient sensory stimuli scaled with the size of evoked pupil dilations. Consistent with a role in sustaining saliency-linked arousal events, pupil responses to sensory stimuli were suppressed with ACC inactivation. Finally, our results comparing arousal-related ACC and norepinephrinergic LC neuron activity support a role for the LC in initiation of arousal events which are modulated in real time by the ACC. Collectively, our experiments identify the ACC as a key cortical site for sustaining momentary increases in arousal and provide the foundation for understanding cortical-subcortical dynamics underlying the modulation of arousal states.

## Main

Fluctuations in waking global arousal are key determinants of spontaneous cortical activity and modulate sensory and task driven responses^1–8^. Global arousal has been measured in these recent studies predominantly as internally driven changes in pupil size, facial movements, and locomotion in the absence of changes in external (i.e., environmental) stimuli. In addition, pupil-linked arousal is associated with other physiological markers of sympathetic tone like heart rate and galvanic skin conductance response^9,10^. Cognitive and emotional behaviors are intimately linked with context-dependent modulation of physiological indicators of arousal^11,12^. Hence, prominent arousal modulation of cortical activity may represent a fundamental mechanism for coordinating central and bodily processes important for behavioral control^13,14^.

Substantial evidence shows that several subcortical structures regulate arousal, including neuromodulatory nuclei that provide widespread cortical outputs like the norepinephrine releasing locus coeruleus (LC) and cholinergic basal forebrain^15–21^. State-dependence of cortical activity is thought to arise in part from a confluence of diverse neuromodulatory influences and other long-range inputs^20,22,23^. Although mechanisms mediating state-dependent modulation of cortical activity have received intense scrutiny, how cortical activity itself modulates arousal and the behavioral state remains an outstanding question.

The anterior cingulate cortex (ACC) subdivision of the prefrontal cortex (PFC) is well-positioned to modulate arousal and physiological states more broadly^24–26^. It provides direct and indirect outputs to multiple neuromodulatory nuclei that coordinate arousal^15,27,28^ and to other subcortical nuclei regulating autonomic function like the periaqueductal gray and the posterior hypothalamus^29^. Electrical stimulation of the human ACC increases the heart rate and engages the skin conductance response^30^. Neuroimaging studies in humans during the Stroop task show that ACC activity correlates with trial-by-trial variations in pupil-linked autonomic arousal^31^. Humans with ACC damage mostly recover their cognitive abilities but show a lasting deficit in task-driven modulation of arousal states^26^. Single-unit recordings in monkeys show increased activity in a small subset of ACC neurons before spontaneous increases in pupil-linked global arousal^32^. Moreover, ACC lesions in monkeys blunt anticipatory pupil arousal responses preceding rewards^33^, further supporting a role for this region in arousal modulation.

Although much evidence supports a role for the ACC in task-driven arousal modulation, how the ACC modulates spontaneous fluctuations in global arousal is not known. Moreover, how arousal-related ACC activity compares to the activity of subcortical neuromodulatory nuclei requires further examination to determine whether the ACC plays a unique role in arousal modulation. We used pupil size, facial movement, heart rate, and locomotion as non-invasive measures of arousal and behavioral state in mice. To test the role of the ACC in arousal modulation, we designed a system for closed loop ACC inhibition based on real-time tracking of the pupil. Optogenetic ACC inactivation after initiation of pupil dilations suppressed ongoing arousal events. In agreement with a role for the ACC in arousal modulation, bulk ACC calcium activity recorded via fiber photometry scaled with pupil size and amplitude of facial movements with a delay after the onset of arousal events and independently of locomotion. Arousal changes evoked by salient sensory stimuli scaled with ACC activity and were suppressed by ACC inactivation, suggesting that the ACC modulates both global and saliency-linked arousal responses. Finally, comparing arousal-related ACC and norepinephrine locus coeruleus (LC-NE) activity suggested that LC-NE activity triggers transient increases in arousal while ACC activity plays a role in sustaining these events. Together, our experiments show that ACC activity modulates the intensity of global and saliency-linked arousal states and establish this PFC region as a cortical site for arousal modulation.

## Results

### Open and closed loop optogenetic ACC inactivation decreases arousal

We used open loop (Extended Data Fig. 1) and closed loop optogenetics (Fig. 1 and Extended Data Fig. 2) to test the role of ACC activity in modulation of global arousal, defined here as spontaneous fluctuations in pupil size occurring in the absence of any presented stimuli. We bilaterally injected VGAT-Cre mice with AAV5-Flx-ChR2 and implanted a fiber optic cannula above the injection sites to inhibit ACC activity via photostimulation of GABAergic neurons^34^ (Extended Data Fig. 1A, B). Control mice were injected with AAV5-CaMKII-mCherry. We recorded the pupil of head-fixed mice with an infrared camera and used DeepLabCut^35^ to track eight key points around the perimeter of the pupil. Pupil size was quantified as the area of an ellipse fit to these key points (Extended Data Fig. 1C). ACC inactivation (20 Hz, 5s duration) decreased the average pupil size while there was no effect of photostimulation in mCherry expressing mice (Extended Data Fig. 1D, E). The effect of ACC inactivation was dependent on the pupil size at the time of photostimulation. Inactivation produced a larger effect when baseline pupil size was large (Extended Data Fig. 1F), suggesting that reducing ACC activity during ongoing pupil dilation events curtails momentary increases in arousal. Furthermore, the effect of ACC inactivation on the pupil size was dependent on the duration of photostimulation (0.5-5s), but we did not detect a significant effect of photostimulation frequency on the decrease in pupil size (Extended Data Fig. 1G, H). Together, these results show that ACC activity modulates spontaneous changes in arousal.

**Figure 1.**
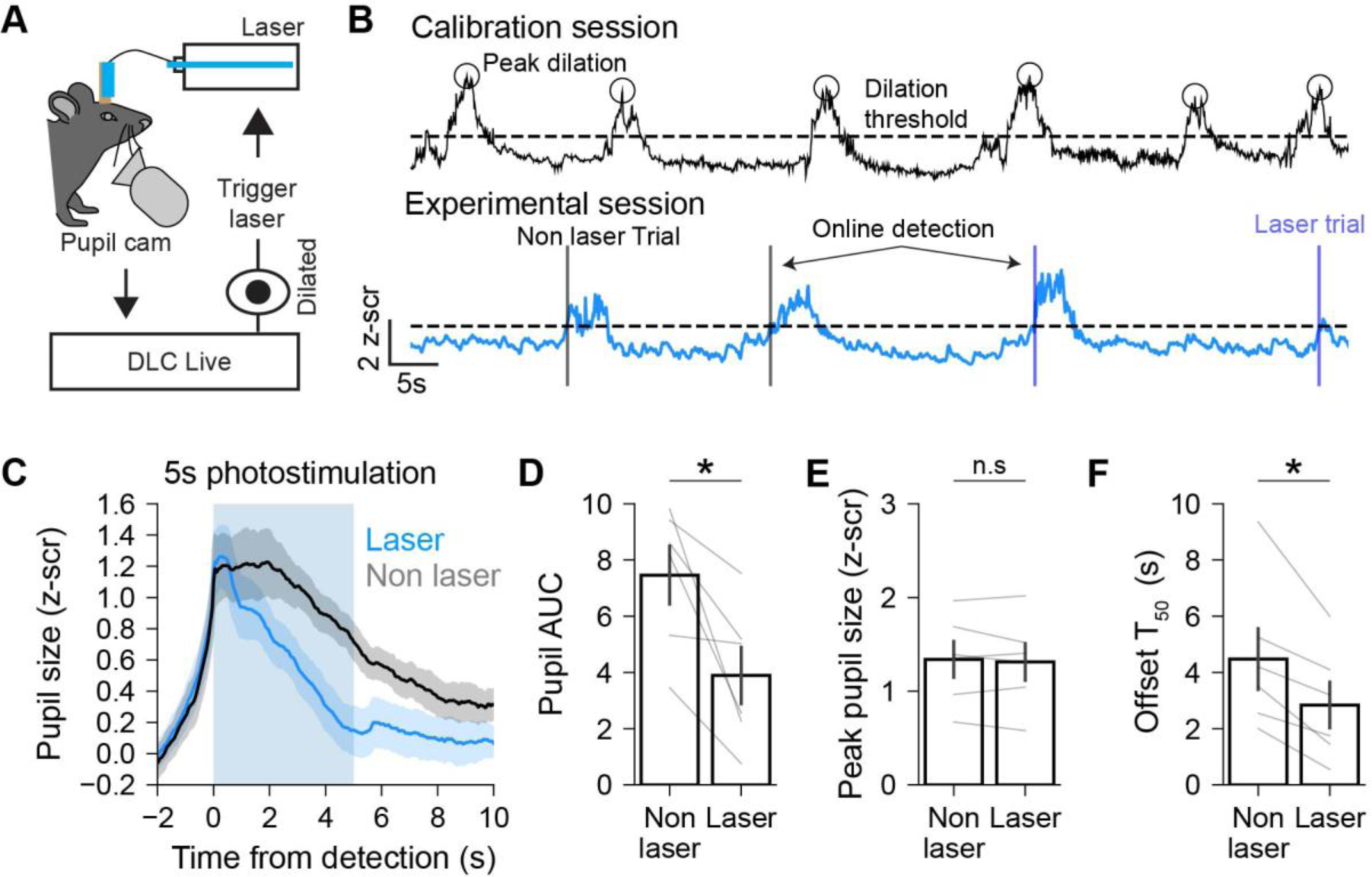
Closed loop ACC inactivation curtails ongoing pupil dilation events. **(A)** Schematic illustrating the experimental setup used to perform closed loop optogenetic stimulation based on real time pupil tracking. **(B)** *Top*, example trace from a calibration session. Circles show pupil dilation peaks. Horizontal dotted line shows the threshold value taken as 25% of the average peak height. *Bottom*, example trace from an experimental session. Vertical lines show online dilation detections for non-laser (black) and laser trials (blue). Horizontal line shows same threshold value as top. **(C)** Pupil size aligned to the time of dilation detection for non-laser and laser trials. Blue shading shows photostimulation time for laser trials. **(D)** Area under the curve (AUC) for non-laser and laser trials (n = 6 mice, p = 0.03, T = 0; Wilcoxon signed-rank test). **(E)** Peak pupil size reached during laser and non-laser trials (n = 6 mice, p = 0.44, T = 6; Wilcoxon signed-rank test). **(F)** Time taken for pupil to decline to 50% of peak value (n = 6 mice, p = 0.03, T = 0; Wilcoxon signed-rank test). For all figures, the Wilcoxon signed-rank test T statistic refers to the number of ranks of differences that are greater than or less than 0 (depending on which is smaller). All error bars are standard error of the mean.

The above results suggest that ACC activity during ongoing pupil dilations is important for arousal modulation. To rigorously evaluate this idea, we developed an experimental paradigm to perform closed loop optogenetic inactivation of the ACC during ongoing pupil dilations (Fig. 1A). We quantified the pupil size in real-time using DeepLabCut-Live^36^ and photostimulated ChR2 expressing GABAergic neurons (20 Hz, 5s duration) on a random 50% of trials (laser) when pupil size was larger than a dilation threshold determined during a preceding calibration session (see Methods; Fig. 1B). The remaining trials were not photostimulated and treated as control (non-laser). We note that in our implementation of this method, photostimulation occurs after the onset of pupil dilations. As a test of the reliability of online detection, we compared the actual pupil size at the time of online detection to the expected size based on the dilation threshold determined during the calibration session. The actual pupil size was slightly higher than the expected size (Extended Data Fig. 2A), which largely reflects the condition that online detected events surpass the threshold determined during calibration. Importantly, there was no difference in the pupil size at the time of online detection for laser and non-laser trials (Extended Data Fig. 2B).

We aligned pupil size to the time of online detection and compared the pupil AUC (area under the curve) between laser and non-laser trials. ACC inactivation significantly reduced the pupil AUC compared to non-laser trials (Fig. 1C, D). Although the average peak pupil size was similar between laser and non-laser trials (Fig. 1E), ACC inactivation led to a faster constriction of the pupil following the peak, as evidenced by a reduction in the time needed for the pupil size to decline to 50% of peak amplitude (Fig. 1F). Repeating this experiment with a shorter duration photostimulation (0.5s) also suppressed ongoing pupil dilation events (Extended Data Fig. 2C-F). There was a decrease in the pupil AUC for laser trials as compared to non-laser trials (Extended Data Fig. 2D), although we could not detect a change in the offset timing of the pupil event with this experiment. These results are unlikely due to virus expression or light delivery in the brain since closed loop optogenetic manipulations in mice injected with AAV-CaMKII-mCherry had no effect on arousal events (Extended Data Fig. 2G-J).

We additionally tested whether the observed suppression of ongoing pupil dilations is specific to inactivation of ACC activity or could be generally observed by inhibiting activity in a control cortical region. We bilaterally injected AAV-Flx-ChR2 into the primary visual cortex (V1) of VGAT-Cre mice and performed the same optogenetic experiments as described above for the ACC. Open loop and closed loop inactivation of V1 activity had no systematic effect on the pupil size (Extended Data Fig. 2K-P). Together, these experiments show that real time ACC activity during pupil dilations is necessary for sustaining ongoing arousal events.

### Simultaneous recording of ACC activity and global arousal

To further determine the relationship between ACC activity and global arousal, we measured bulk ACC calcium activity simultaneously with spontaneous fluctuations in the behavior of head-fixed mice. We injected AAV5-Syn-GCaMP8m into the ACC and implanted a fiber optic to record population level calcium activity. Video recordings of the face were used to quantify both pupil size and facial movement as metrics of global arousal (Extended Data Fig. 3A). We detected individual pupil dilations and quantified event metrics including time of onset, offset, and peak amplitude (dashed lines in Extended Data Fig. 3B; Extended Data Fig. 4; see Methods). Facial movement was defined by taking frame-by-frame differences in mean pixel intensity of an ROI centered around the whisker pad (Extended Data Fig. 3A). In addition, mice were allowed to run voluntarily on a wheel which was used to quantify running speed and determine spontaneous changes in the behavioral state (Extended Data Fig. 3B). Pupil size, facial movements, and running speed were moderately correlated with each other (Extended Data Fig. 3C). Cross-correlation analyses showed that, on average, an initial increase in facial movement was followed by an increase in pupil size and finally by an increase in wheel speed (Extended Data Fig. 3D, E). Applying a moving median filter to smooth the face signal increased the pupil-face correlation while smoothing the pupil signal had no effect (Extended Data Fig. 3F, G), suggesting that pupil and face metrics of arousal partly convey related arousal information but with different temporal dynamics.

To better interpret how ACC activity modulates arousal, we further explored the interrelationship between pupil size, facial movements, and locomotion. Most pupil dilations were short lasting (<5s) and had small peak amplitude (<1 z-score; Extended Data Fig. 3H). Splitting pupil events into quartiles based on the dilation amplitude showed that, on average, large amplitude pupil dilations also had long durations (Extended Data Fig. 3I). Aligning the face signal to pupil dilation onsets showed that both small and large pupil dilations were preceded by facial movements, and the magnitude of facial movements increased progressively with pupil dilation amplitude (Extended Data Fig. 3J). In contrast, only large pupil dilations were associated with locomotion (Extended Data Fig. 3K-N). Like previous findings^37^, these analyses show that while spontaneous fluctuations in arousal can occur in the absence of locomotion, large increases in global arousal are associated with behavioral state shifts.

We tested whether pupil size is a reliable indicator of autonomic arousal by measuring the resting heart rate via pulse oximetry simultaneously with pupil size in a subset of mice (Extended Data Fig. 5A). Spontaneous changes in resting heart rate coincided with pupil dilations (Extended Data Fig. 5B) and a dilated pupil state was associated with higher heart rate (Extended Data Fig. 5C). Splitting pupil events into quartiles based on the dilation amplitude showed that the heart rate increased with pupil size (Extended Data Fig. 5C, D). Hence, pupil size is a reliable readout of autonomic arousal.

### ACC activity scales with the magnitude of arousal events

Our optogenetic results are consistent with the idea that ACC activity is important for sustaining arousal events (Fig. 1, Extended Data Fig. 1 and Extended Data Fig. 2). Next, we investigated how ACC population activity relates to naturally occurring spontaneous fluctuations in global arousal. Since arousal and locomotion are closely linked, we determined how arousal-related ACC activity depends on the locomotion state. We aligned ACC activity to pupil dilation onsets occurring during locomotion and non-movement. Both types of pupil dilations were coincident with increased ACC activity; however, arousal-related activity was higher during locomotion than during non-movement (Extended Data Fig. 6A-C). This effect could reflect larger dilation pupil events that occur during locomotion (Extended Data Fig. 3L). Alternatively, higher levels of ACC activity might reflect the locomotion state itself. We distinguished between these possibilities by comparing arousal-related ACC activity during small and large pupil events occurring during locomotion and non-movement periods (Fig. 2A-D). Small and large dilations were defined using a median split of peak dilation amplitudes observed during each condition within single recording sessions. Compared to small dilation events, large pupil dilations were associated with higher ACC activity during locomotion and non-movement. Importantly, there was no difference in the average locomotion speed for small and large pupil dilations in both conditions (Fig. 2B, D), suggesting that differences in movement speed do not account for increased ACC activity during larger pupil dilations. These findings show that ACC activity tracks the magnitude of arousal events regardless of the locomotion state.

**Figure 2.**
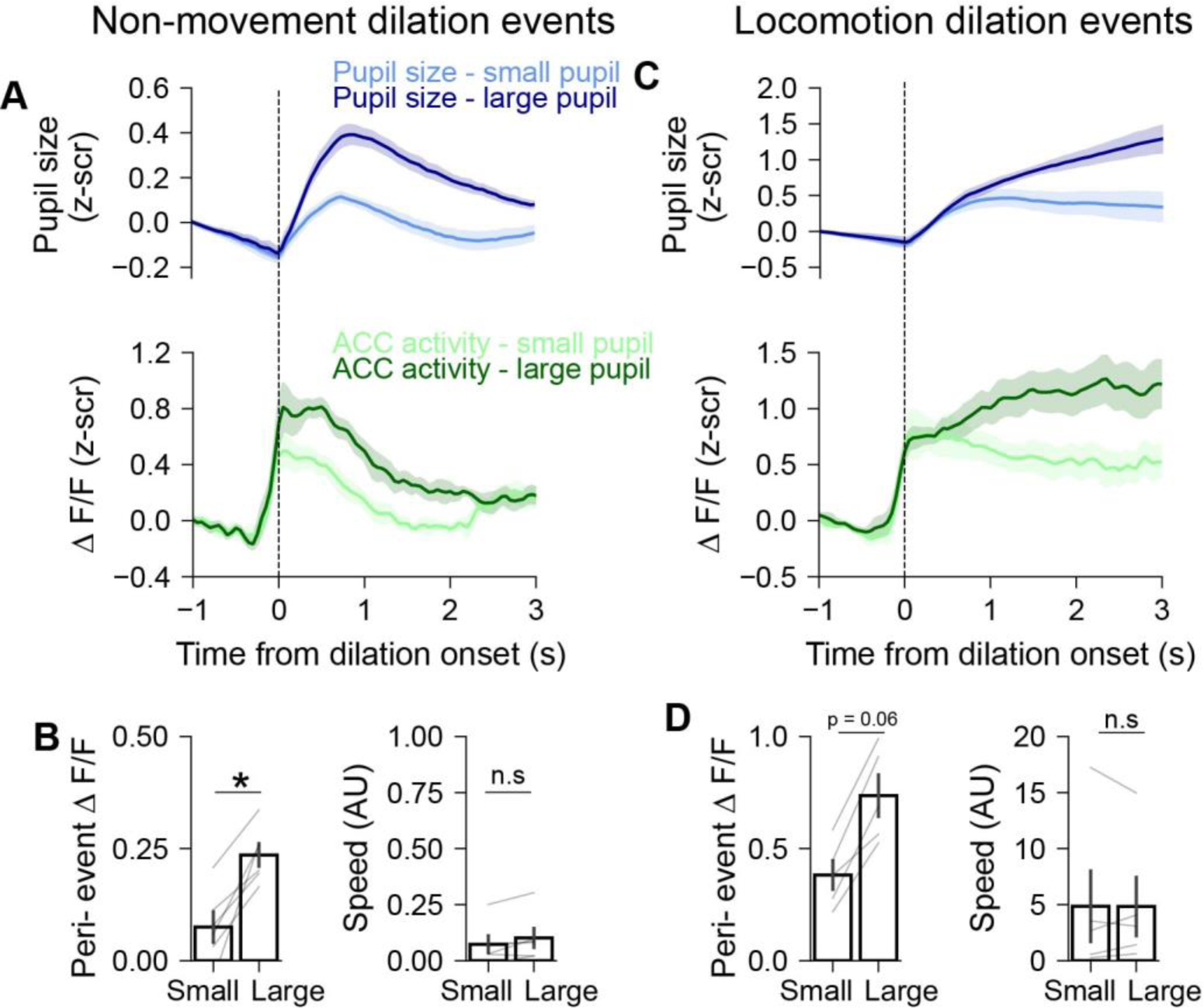
ACC activity scales with the magnitude of arousal events. **(A)** *Top*, pupil events occurring during non-movement aligned to the time of dilation onset. Events are separated into small and large events based on a median split of peak amplitudes. *Bottom*, ACC activity during non-movement small and large events (n = 6 mice). **(B)** ACC activity (left) and mean locomotion speed (right) for small and large pupil dilations during non-movement (activity: n = 6 mice, p = 0.03, T = 0, Wilcoxon signed-rank test; speed: n = 6 mice; p = 0.09, T = 2; Wilcoxon signed-rank test). **(C)** Onset aligned pupil events (top) and ACC activity (bottom) for small and large events during locomotion (n = 6 mice). **(D)** ACC activity (left) and mean locomotion speed (right) for small and large pupil dilations during locomotion (activity: n = 5 mice; p = 0.06, T = 0, Wilcoxon signed-rank test; speed: n = 5 mice; p = 1, T = 7; Wilcoxon signed-rank test). All error bars are standard error of the mean.

As a further test of arousal-related ACC activity, we determined the relationship between ACC activity and facial movements. We aligned ACC activity and facial movements to the onset time of pupil dilation events and separated trials based on small and large facial movements (Extended Data Fig. 6D and 6E). Large facial movements were accompanied by higher ACC activity as compared to small facial movements (Extended Data Fig. 6F). Hence, ACC activity scales with the magnitude of pupil dilations and facial movements. Together with the open and closed loop optogenetic inactivation results (Fig. 1 and Extended Data Fig. 1), these analyses show that the scaling of arousal-related ACC activity contributes to the magnitude of ongoing arousal events.

### Optogenetic activation of ACC neurons increases arousal and locomotion

Our results show that ACC activity correlates with the magnitude of arousal events and inhibiting ACC activity suppresses ongoing pupil events (Fig. 1 and Fig. 2), suggesting that an increase in ACC activity should lead to higher arousal. We tested this by determining the effect of optogenetic ACC activation on arousal and locomotion. We virally expressed CaMKII-ChR2 and implanted a fiber optic in the ACC to deliver light and optogenetically activate ACC neurons (Fig. 3A, B). Optogenetic activation (20 Hz, 5s duration) increased pupil size, wheel running speed and facial movements (Fig. 3C-G). Pupil size and running speed generally increased throughout the duration of photostimulation and decreased after stimulation offset (Fig. 3D, F). However, facial movement peaked shortly after stimulation onset, declined during the photostimulation epoch and showed a second increase after stimulation offset (Fig. 3G). Importantly, photostimulation in mice expressing the mCherry control fluorophore did not change the pupil size (Fig. 3D, E). We found that the effect of photostimulation on arousal metrics was frequency dependent with higher frequencies evoking larger pupil dilations and facial movements as well as higher locomotion speeds (Extended Data Fig. 7A-D). The effect of stimulation on pupil size and running speed was also duration dependent with longer duration photostimulation leading to larger and longer lasting increases (Extended Data Fig. 7E-G). In contrast, the effect of stimulation on facial movement was not duration dependent (Extended Data Fig. 7H). In a subset of mice, we also measured the resting heart rate and found that optogenetic ACC activation increased the heart rate (Extended Data Fig. 7I, J). These results demonstrate that seconds long ACC activation increases diverse metrics of arousal and leads to behavioral state shifts.

**Figure 3.**
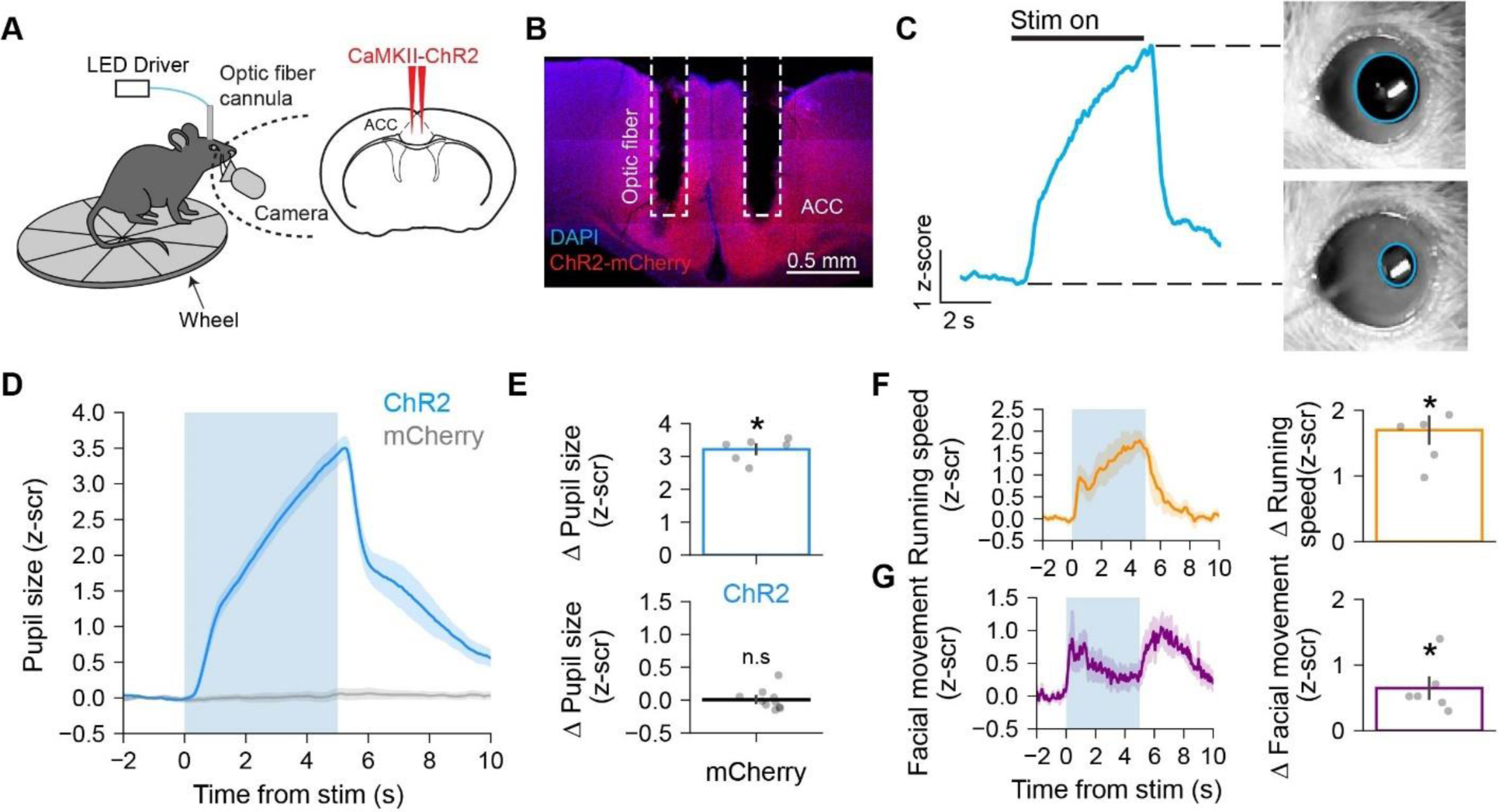
Optogenetic ACC activation increases arousal and locomotion. **(A)** Schematic illustrating the experimental setup for measuring arousal and locomotion during optogenetic activation of CaMII-ChR2 expressing ACC neurons. **(B)** Coronal section showing CaMKII-ChR2-mCherry expression and fiber optic placement in the ACC. **(C)** Single trial example for the effect of optogenetic activation on pupil size. Video images show the pupil at the beginning and the end of 5s long photostimulation. **(D)** Pupil size aligned to photostimulation onset for mice expressing ChR2-mCherry or mCherry (n = 6 and 10 mice for ChR2 and mCherry conditions, respectively). Blue shading shows the photostimulation window. **(E)** *Top*, Photostimulation evoked change in pupil size for ChR2 mice (n = 6 mice, p = 0.03, T = 0; Wilcoxon signed-rank test against zero). *Bottom*, Same as top but for mCherry controls (n = 10 mice, p = 0.85, T = 25 Wilcoxon signed-rank test against zero). **(F)** *Left*, Running speed aligned to photostimulation onset. *Right*, photostimulation evoked change in running speed (n = 6 mice, p = 0.03, T = 0; Wilcoxon signed-rank test against 0). **(G)** *Left*, Facial movement aligned to photostimulation onset. *Right*, photostimulation evoked change in facial movement (n = 6 mice, p = 0.03, T = 0; Wilcoxon signed-rank test against 0).

### Short duration ACC activation increases arousal independently of locomotion

Optogenetic ACC activation evoked increases in both arousal and locomotion. Since locomotion is closely linked with high arousal levels (Extended Data Fig. 3L, M), it is possible that ACC activation primarily triggers locomotion and increased arousal is a secondary effect. We tested whether ACC activation can increase arousal independently of locomotion by delivering a single 10ms pulse of optogenetic stimulation. In this open loop experiment, photostimulation could occur during periods of wheel quiescence or locomotion. We concatenated data across mice and split trials based on locomotion state depending on the average wheel speed around photostimulation (window: −0.5s to 2s relative to stimulation). Short duration ACC activation increased pupil size and facial movement during non-locomotion trials (Fig. 4A-C), demonstrating that ACC activity can modulate arousal independently of locomotion. ACC activation during locomotion also increased pupil and facial movement metrics of global arousal (Fig. 4D-F). The change in pupil size with ACC activation was greater during locomotion than during quiescence (n = 599 non-locomotion trials, 424 locomotion trials, p <0.00001, U = 63700; Mann-Whitney U rank test), suggesting that brief elevation of ACC activity during the high arousal state associated with locomotion produces a stronger effect. ACC activation in this experiment had no significant effect on wheel speed, ruling out the possibility that observed increase in pupil size following ACC activation were driven by changes in locomotion. These findings show that minimal ACC activation increases arousal independently of locomotion, mirroring our observations with ACC fiber photometry recordings (Fig. 2).

**Figure 4.**
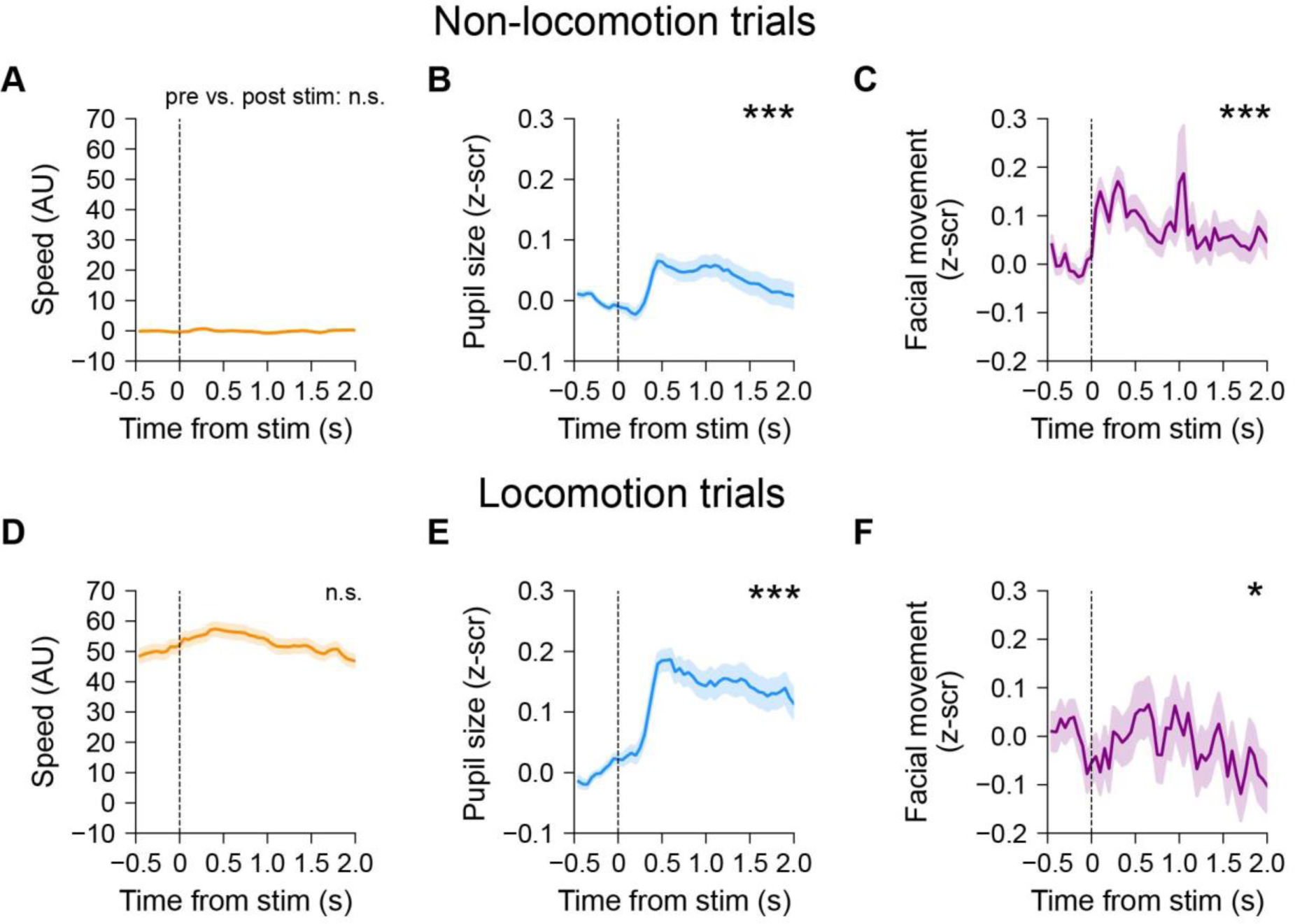
Short duration ACC activation increases arousal independently of locomotion. **(A)** Running speed aligned to photostimulation onset for non-locomotion trials (n = 599 trials, p = 0.95, T = 69879; Wilcoxon signed-rank test). **(B**) Same as A but for pupil size (n = 599 trials, p = 0.0005, T = 74984; Wilcoxon signed-rank test). **(C)** Same as A but for facial movement (n = 599 trials, p <0.00001, T = 63700; Wilcoxon signed-rank test). **(D)** Running speed aligned to photostimulation onset for locomotion trials (n = 424 trials, p = 0.16, T = 41483; Wilcoxon signed-rank test). **(E**) Same as D but for pupil size (n = 424 trials, p <0.00001, T = 26409; Wilcoxon signed-rank test). **(F)** Same as D but for facial movement (n = 424 trials, p = 0.03, T = 39517; Wilcoxon signed-rank test). Asterisks signify significant difference between the average value in pre-stim (−0.5s to 0s) and post-stim (0s to 2s) windows. All error bars are standard error of the mean.

### ACC activity modulates arousal associated with salient stimuli

Arousal levels change in response to salient stimuli^12^ in addition to fluctuating spontaneously. Given the role of the ACC in modulating spontaneous changes in global arousal, we determined how the ACC contributes to stimulus evoked pupil dilations. We measured ACC population activity and pupil size during auditory stimulation consisting of pure tones at 3 different frequencies (Fig. 5A, B). On average, auditory stimulation increased pupil size and ACC activity (Fig. 5C). There was significant variability in the pupil response to the tone (Fig. 5D). There was no dependence of pupil or ACC responses on tone frequency; moreover, tone evoked pupil dilations and ACC responses were similar across the recording session (Extended Data Fig. 8). These results suggest that tone frequency-specific effects or habituation are unlikely contributors to the observed variability. To determine if the observed variability in the magnitude of the elicited pupil response was related to ACC activity, we split tone-evoked pupil dilations into quartiles. Comparing ACC activity across quartiles showed that higher ACC responses during auditory stimulation were associated with larger arousal events (Fig. 5E-G). These results show that ACC activity correlates with the magnitude of saliency-linked arousal responses.

**Figure 5.**
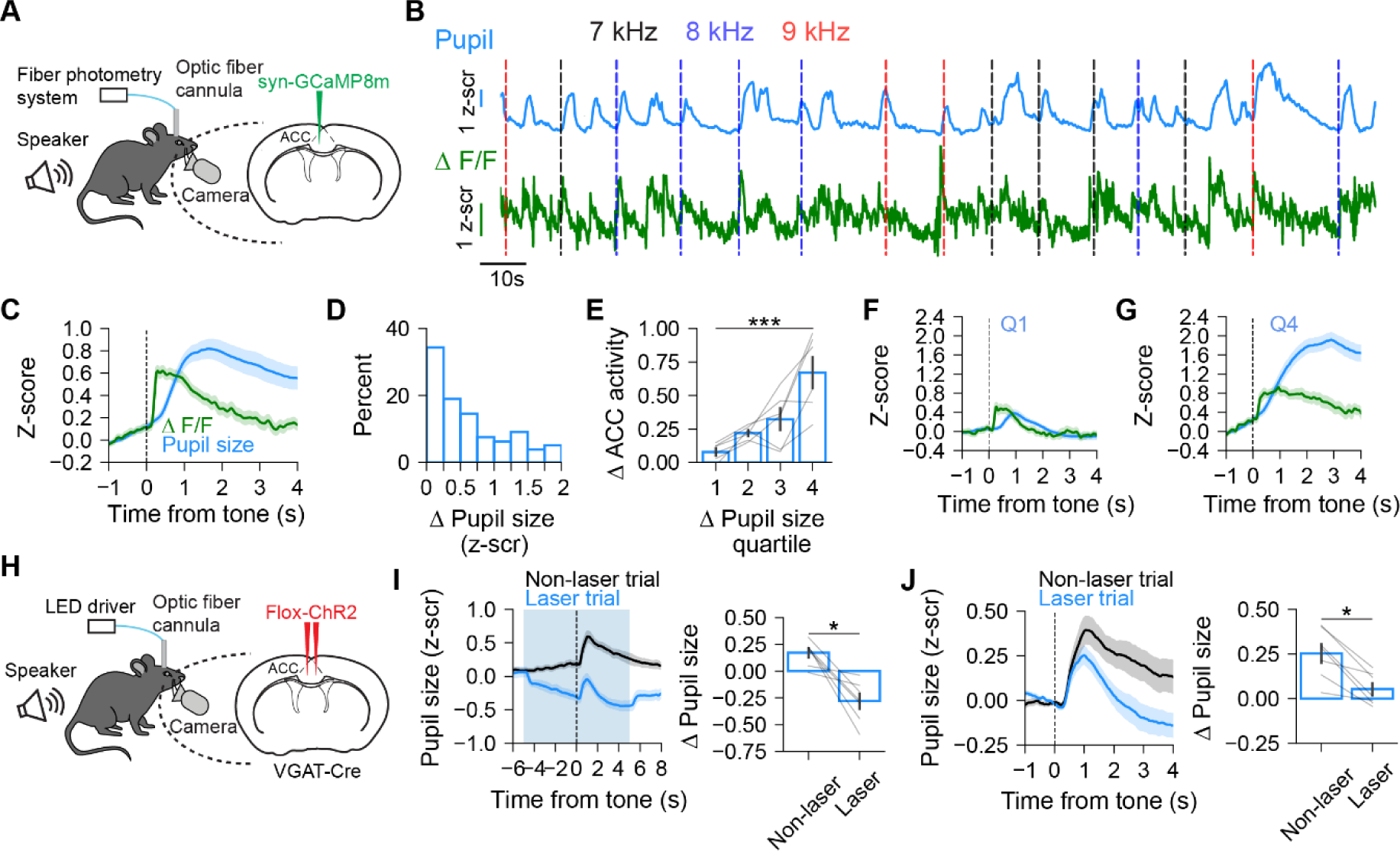
ACC activity modulates arousal associated with salient stimuli. **(A)** Schematic illustrating the experimental setup used to simultaneously record pupil size and population level ACC activity during presentation of auditory tones. **(B**) Example traces from a session showing pupil size and ACC activity during presentation of auditory tones. Vertical lines show tone onsets. **(C)** ACC activity and pupil size aligned to tone presentation (n = 6 mice). **(D)** Histogram showing the distribution of average changes in pupil size in response to tone presentation (n = 6 mice). **(E)** Changes in ACC activity in response to tone presentation across quartiles of pupil size changes (n = 6 mice; p = 0.002, H = 14.98; Kruskal-Wallis test). **(F)** ACC activity and pupil size aligned to onset of tone presentation for trials in Q1 (first quartile) (n = 6 mice). **(G)** Same as F, but for Q4 (fourth quartile) (n = 6 mice). **(H)** Schematic illustrating the experimental setup used to record pupil size and optogenetically inhibit ACC activity during presentation of auditory tones. **(I)** *Left*, Pupil size aligned to tone presentation for non-laser and laser trials. *Right*, Change in pupil size for non-laser and laser trials during entirety of laser stimulation (n = 7 mice, p = 0.02, T = 0; Wilcoxon signed-rank test). **(J)** *Left*, Pupil size aligned to tone presentation for non-laser and laser trials (baseline corrected using 1s preceding tone). *Right*, Change in pupil size for non-laser and laser trials in the 3s following tone presentation (n = 7 mice, p = 0.02, T = 0; Wilcoxon signed-rank test). All error bars are standard error of the mean.

We directly tested how ACC activity contributes to tone-evoked arousal responses by inactivating the ACC using ChR2-expressing VGAT-Cre mice (Fig. 5H). On each laser trial, photostimulation began five seconds before tone onset and lasted for ten seconds. As we observed before (Extended Data Fig. 1), ACC inactivation decreased global arousal, which was marked by a decrease in the pupil size in trial-averaged traces (Fig. 5I). Because of this, baseline pupil size was lower on laser trials as compared to non-laser trials (Fig. 5I). To examine the effect of ACC inactivation on sensory evoked arousal specifically, we used a 1s period preceding tone onset to baseline correct all trials. This showed that while ACC inactivation did not abolish the sensory evoked arousal response, it decreased the magnitude of the response (Fig. 5J). This result further supports the idea that ACC activity is important for sustaining elevated arousal states elicited by salient stimuli.

### ACC and LC show unique patterns of arousal-related activity

Inactivating the ACC decreased tone-evoked pupil dilations but did not entirely inhibit the arousal response (Fig. 5J). Moreover, in closed loop experiments showing suppression of ongoing arousal events with ACC inactivation (Fig. 1), we could only inactivate the ACC after an initial increase in pupil size. These findings suggest that under naturalistic conditions, ACC activity is important for sustaining increases in arousal rather than initiating arousal events, for which other brain areas may be more important. We tested this idea by recording the bulk calcium activity of norepinephrine releasing LC (LC-NE) neurons and comparing arousal-related LC-NE activity with ACC arousal responses. We injected AAV-Flx-GCaMP8 into the LC of DBH-Cre animals and implanted a fiber optic above the injection site for photometry recordings (Fig. 6A). ACC activity showed higher correlation with pupil size than LC-NE activity (Fig. 6B), an effect possibly driven by the observed scaling of ACC responses with pupil dilation amplitude (Fig. 2). Conversely, facial movement showed a higher correlation with LC-NE activity than ACC activity (Fig. 6B). To directly examine the temporal relationship between ACC/LC activity and arousal events, we aligned neuronal activity to the onset time of pupil dilations (Fig. 6C). Both ACC and LC-NE activity preceded pupil dilations. However, LC-NE activity increased earlier and faster than ACC activity relative to pupil dilation onset (Fig. 6C-E). ACC activity lagged facial movements, a faster readout of arousal than pupil (Extended Data Figs. 3F and 3G), while LC activity increased near simultaneously with facial movements (Fig. 6C). Hence, LC activity increases coincident with the onset of arousal while ACC activity increases with a delay after initiation of the arousal event.

**Figure 6.**
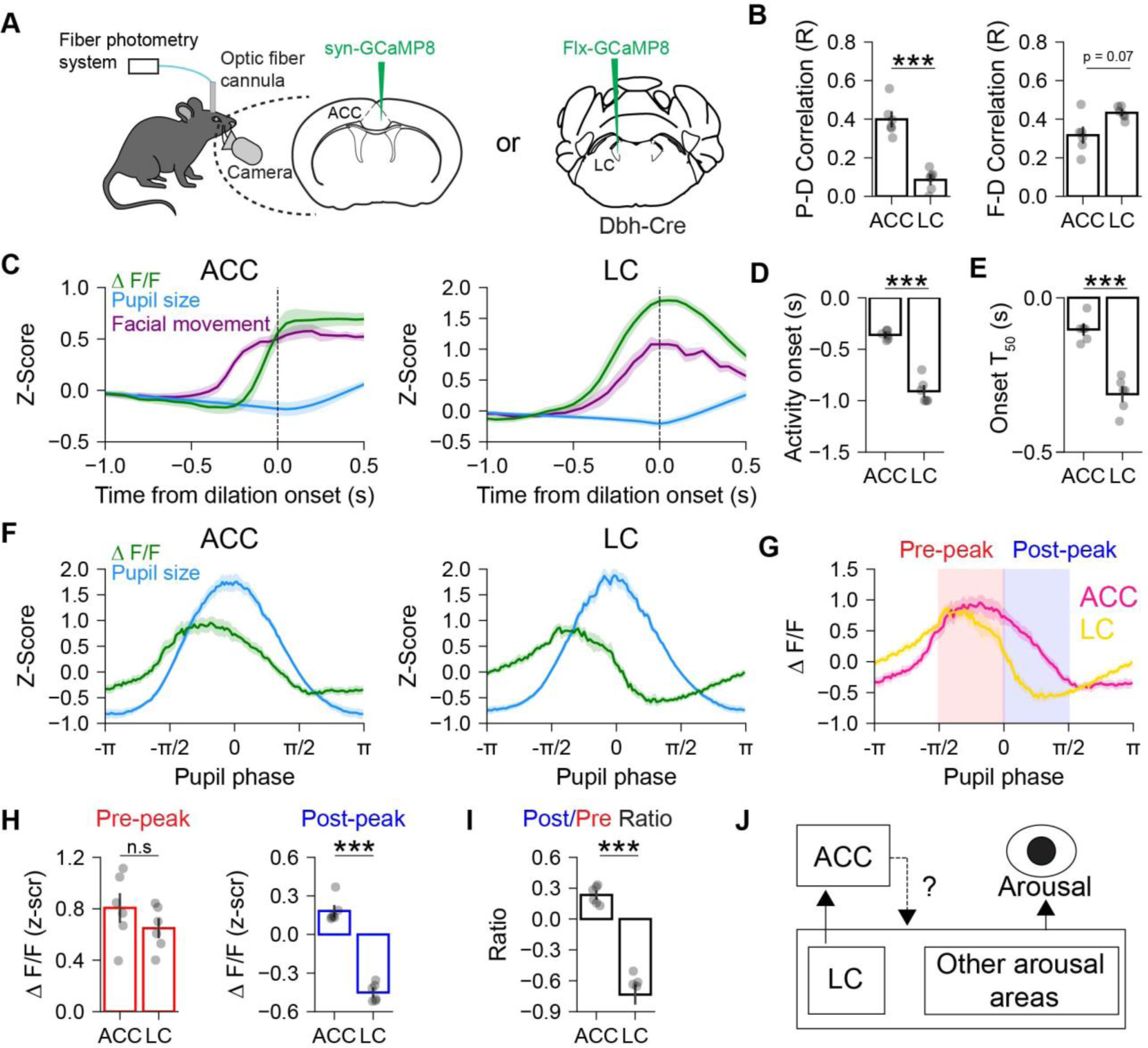
ACC and LC show unique patterns of arousal-related activity. **(A)** Schematic illustrating the experimental setup used to record arousal and measure bulk calcium signals from ACC or LC-NE neurons. **(B)** *Left,* Pearson’s correlation between pupil size and activity in ACC or LC (n = 6 LC mice, 6 ACC mice, p = 0.002, U = 0; Mann-Whitney U rank test). *Right,* same as Left but with facial movement (n = 6 LC mice, 6 ACC mice, p = 0.07, U = 30; Mann-Whitney U rank test). **(C)** *Left,* traces of ACC activity, facial movement, and pupil size aligned to the onset of pupil dilation (n = 6 mice). *Right,* same as left but for LC-NE activity (n = 6 mice). **(D)** Onset of activity relative to the onset of pupil dilation for ACC or LC (n = 6 LC mice, 6 ACC mice, p = 0.005, U = 0; Mann-Whitney U rank test) **(E)** Time to reach 50% of peak activity relative to onset of pupil dilation in ACC or LC (n = 6 LC mice, 6 ACC mice, p = 0.005, U = 0; Mann-Whitney U rank test). **(F)** *Left,* ACC activity and pupil size as a function of pupil phase. *Right*, same as Left but for LC-NE activity. **(G)** ACC and LC-NE activity as a function of pupil phase. Red shaded rectangle represents the pre-peak (-π/2 to 0) window and blue shaded rectangle represents the post-peak (0 to π/2) window (n = 6 mice for ACC and LC respectively). **(H)** *Left,* Average activity during the pre-peak window for ACC or LC (n = 6 LC mice, 6 ACC mice, p = 0.31, U = 11; Mann-Whitney U rank test). *Right*, same as Left but for post-peak window (n = 6 LC mice, 6 ACC mice, p = 0.002, U = 0; Mann-Whitney U rank test) **(I)** Post-peak/pre-peak ratio for ACC or LC (n = 6 LC mice, 6 ACC mice, p = 0.002, U = 0; Mann-Whitney U rank test). **(J)** Model illustrating a possible circuit for ACC interactions with the LC and other arousal-related regions to modulate arousal states.

To better understand ACC and LC-NE activity relative to spontaneous fluctuations in pupil size, we aligned activity to a canonical cycle of pupil dilation and constriction determined using the Hilbert transform^37^ (Fig.6F, G). ACC and LC-NE activity both showed pupil phase-linked fluctuations. Activity in both regions increased around the peak of dilation and decreased during constriction, although LC-NE activity started declining earlier than ACC activity (Fig.6F, 6G). Comparing pupil phase aligned responses showed similar LC-NE and ACC activity preceding peak dilation (Fig. 6H). However, we observed higher ACC activity during the post-peak phase of the dilation compared to LC-NE activity (Fig. 6H). We quantified the ratio of post-peak to pre-peak activity as an index for relative activity during these phases and found that ACC had a higher ratio than LC (Fig. 6I). These findings suggest that compared to LC activity, ACC activity is well placed to modulate the amplitude of already initiated pupil dilation events.

Given the temporal difference in arousal-related ACC and LC-NE activity, we next compared how activity in these regions scales with the magnitude of pupil dilation events. We aligned activity to the time of pupil dilation onset and compared pre-onset and peri-event responses across quartiles of pupil dilation amplitude (Extended Data Fig. 9). Examining the average pupil-aligned responses showed that LC activity started declining around the time of the peak pupil dilation (Extended Data Fig. 9A). This was unlike ACC activity which remained elevated, especially for larger amplitude events (Extended Data Fig. 9C). There was no relationship between pre-onset ACC activity and peak pupil amplitude (Extended Data Fig. 9A and 9B). However, the average level of ACC activity quantified during the pupil event (i.e., from pupil dilation onset to offset; Extended Data Fig. 4) increased with dilation amplitude (Extended Data Fig. 9A, B). These results support our earlier findings that ACC activity scales with the magnitude of arousal events (Fig. 2). We found no systematic relationship between pre-onset LC-NE activity and peak pupil amplitude (Extended Data Fig. 9C, D). In contrast to the ACC, arousal-related LC-NE activity did not scale across the range of observed pupil dilation amplitudes (Extended Data Fig. 9D). Quantifying the difference in activity at the peak of the dilation event from activity at the onset of the event showed that ACC activity changed only slightly following the dilation, but LC-NE activity was drastically reduced (Extended Data Fig. 9E).

In combination with our temporal analysis of arousal-related ACC and LC-NE activity (Fig. 6), results from closed loop optogenetic ACC inactivation experiments, and previous studies^15,38^, these results support a role for the LC in initiation of arousal events which are further modulated in real time by ongoing levels of ACC activity.

## Discussion

There is a growing appreciation for widespread arousal state-dependent modulation of cortical activity. Our experiments demonstrate that in addition to being modulated by arousal, cortical activity in the ACC itself regulates spontaneous and saliency-linked arousal states. Using a closed loop optogenetic design, we show that real-time inhibition of ACC activity during pupil dilations suppresses ongoing arousal events (Fig. 1). Fiber photometry recordings showed that bulk ACC calcium activity scales with the magnitude of arousal events independently of locomotion (Fig. 2). Long duration optogenetic ACC activation drives robust increases in arousal which lead to behavioral state shifts as evidenced by increased locomotion (Fig. 3). However, short duration ACC activation increases arousal without an effect on locomotion (Fig. 4). This demonstrates that ACC modulates arousal independently of behavioral state shifts and increased locomotion, when observed, is a secondary effect to arousal. We found that ACC activity also modulates saliency-linked arousal. ACC activity scales with the amplitude of auditory tone evoked pupil dilations, which are suppressed by optogenetic ACC inhibition (Fig. 5). Comparing arousal-related ACC and LC-NE activity suggests that LC-NE activity may trigger arousal events which are modulated in real time by ACC activity (Fig. 6).

We used pupil size, facial movement, heart rate, and running speed as non-invasive readouts of arousal level and behavioral state. The correlation between pupil size and heart rate (Extended Data Fig. 5) shows a close correspondence between physiological and video-based metrics of arousal. Like previous work^37^, we found that although locomotion is associated with large pupil dilations, spontaneous fluctuations in pupil size and facial movement also occur during periods of non-movement. These findings indicate that arousal and locomotion are related but separate states, an idea further supported by studies showing differential modulation of visual cortical activity by pupil size and running^4^. Pupil size, heart rate, and facial movement may represent observable features of a latent arousal variable. However, to what extent each of these variables represents distinct and overlapping information is unclear. Smoothing the facial movement signal increases its correlation with pupil size (Extended Data Fig. 3G), suggesting that it partly resembles the pupil variable but operates with faster temporal dynamics. The difference in temporal dynamics may be in part due to the muscles involved in controlling each response. One practical consequence is that the slower nature of the pupil signal may allow it to integrate internal state and other information over longer time periods than facial movement.

We took advantage of this temporal feature in this study to design a closed loop optogenetic system based on real-time pupil tracking. In our implementation of this technique, optogenetic manipulation occurs after the pupil dilation event has already started. Yet, we observed that ACC inactivation reliably curtailed ongoing pupil dilation events (Fig. 1). Consistent with our fiber photometry results (Fig. 2 and Extended Data Fig. 9), this finding shows that ACC activity after the onset of arousal events is important for arousal modulation. More broadly, our results show that neuronal manipulations based on real time measurement of behavioral signals can give valuable insights on the functional role of specific brain circuits.

The activity of LC-NE neurons has long been implicated in arousal modulation ^11,15,16,20,32,38^. Comparing bulk calcium activity of ACC and LC-NE neurons showed that LC-NE activity increases near simultaneously with the onset of facial movements while ACC activity increases after a delay, suggesting that ACC activity lags LC activation (Fig. 6). This sequence of events suggests that under typical conditions, ACC activity does not contribute to the initiation of arousal events. In contrast to LC-NE activity, ACC activity scales with the size of pupil dilations, suggesting that ACC activity modulates ongoing arousal events (Extended Data Fig. 9). This relationship between ACC activity and arousal event amplitude was observed during both non-movement and locomotion (Fig. 2), demonstrating that ACC modulates arousal independently of the behavioral state. One possibility is that ACC activity before pupil dilation onset reflects neuromodulatory inputs from LC-NE neurons and other sources. Indeed, several studies have shown that increases in arousal are associated with widespread increases in both cholinergic and noradrenergic activity across the cortex^22,23^. In this interpretation, neuromodulatory inputs trigger the initial increase in ACC activity during arousal events, with subsequent arousal-related modulation resulting from integration within the ACC (Fig. 6J). This idea is also consistent with previous work showing only moderate coupling between moment-by-moment variations in LC activity and pupil-linked arousal^39^. Our work further suggests that variability in pupil-linked arousal partly reflects ACC activity. Future experiments are needed to determine how neuromodulatory inputs contribute to ACC modulation of arousal and how the ACC regulates activity in arousal-related neuromodulatory nuclei.

While the above results show that ACC activity is important for modulating ongoing arousal events, we also found that direct optogenetic ACC activation triggers de novo pupil dilations and facial movements (Figs. 3 and 4). Activation of numerous subcortical brain areas including the lateral hypothalamus, nucleus incertus, amygdala, ventral midline thalamus and PAG is associated with increased autonomic arousal^18,19,21,40^. The ACC sends direct and indirect projections to many of these subcortical regions^29^ which when recruited by optogenetic activation could result in pupil dilation. Regardless of the exact circuit mechanisms underpinning these results, our fiber photometry experiments show that average ACC activity increases after facial movement. This finding suggests that under naturalistic conditions arousal events are not initiated by ACC activity. At the same time, these experiments also demonstrate the possibility of triggering arousal events directly via increased ACC activity, a pathway which may be recruited under pathological conditions. Further experiments are needed to clarify the exact output projections from the ACC that can modulate global and saliency-linked arousal.

In recent years, increased attention has been placed on the influence of arousal and behavioral state on cortical activity. Multiple studies have shown that variability in cortical activity is explained to a high degree by spontaneous fluctuations in arousal^1–8^. Our results imply the possibility that the ACC can exert large-scale effects on cortical processing by modulating arousal.

## Extended data figures

**Extended Data Fig. 1:**
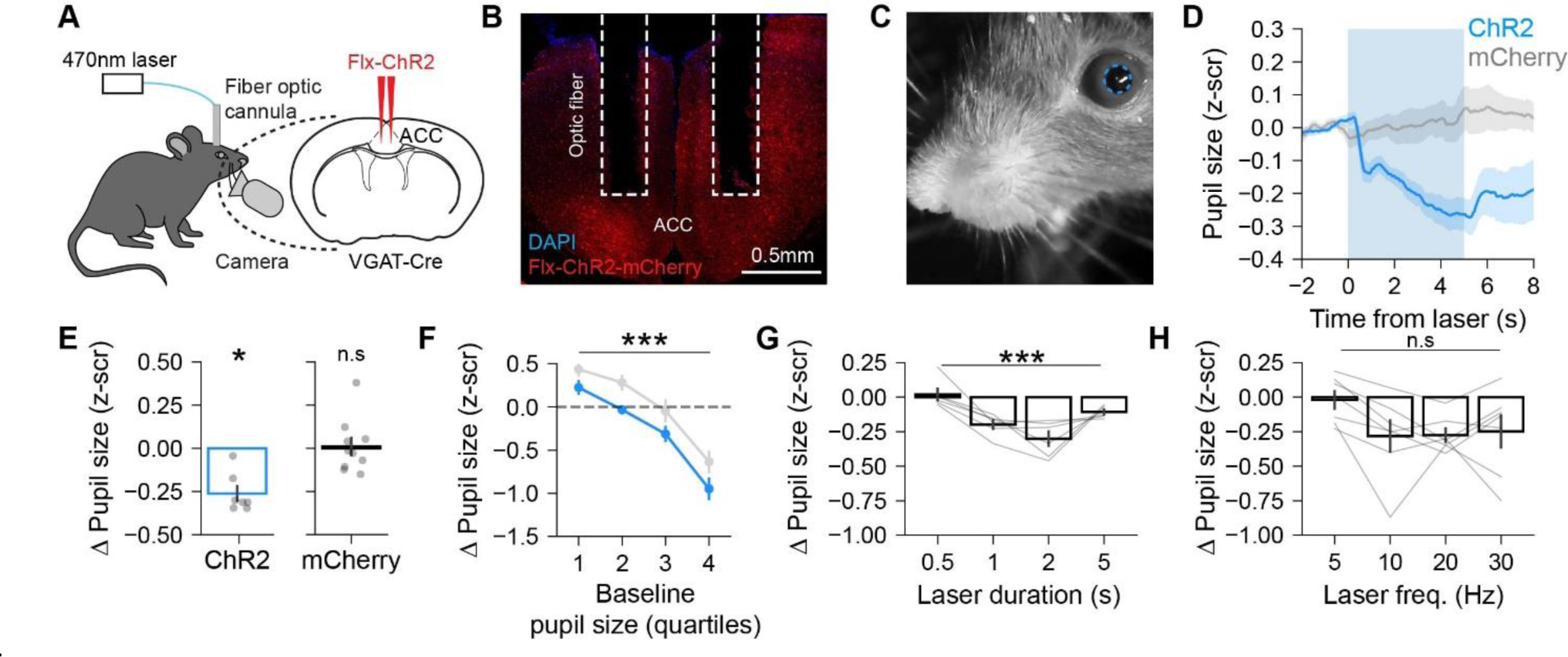
Open loop optogenetic ACC inactivation decreases arousal. **(A)** Schematic illustrating the experimental setup used to record pupil size and optogenetically inhibit ACC activity. Inhibition of ACC neurons was done by activating GABAergic neurons in VGAT-Cre mice injected with Flx-Chr2. **(B)** Coronal section of the ACC showing Flx-ChR2-mCherry expression and fiber optic placement in the ACC. **(C)** Example video frame of the face used for quantifying pupil size (dashed blue line). **(D)** Pupil size aligned to photostimulation onset for experimental mice expressing ChR2-mCherry or control mice expressing only mCherry (n = 7,10 mice for ChR2 and mCherry conditions respectively). **(E)** *Left*, photostimulation evoked change in pupil size for ChR2 mice (n = 7 mice, p = 0.02, T = 0; Wilcoxon signed-rank test against zero). *Right*, same as left but for mCherry controls (n = 10 mice, p = 0.85, T = 25; Wilcoxon signed-rank test against zero). **(F)** Photostimulation evoked change in pupil size plotted as a function of pre-laser (baseline) pupil size in quartiles for ChR2 and mCherry (n = 7 mice for ChR2 group, 10 mice for mCherry group; main effect of baseline pupil size quartile F (3,45) = [53.27], p = 7.5e-15; main effect of virus group F (1,15) = [15.24], p = 0.001; two-way mixed measures ANOVA). **(G)** Change in pupil size evoked by photostimulation at varying durations (n = 6 mice; p = 0.0003, H = 13.99; Kruskal-Wallis test). **(H)** Change in pupil size evoked by photostimulation across varying frequencies (n = 7 mice; p = 0.09, H = 6.6; Kruskal-Wallis test).

**Extended Data Fig. 2:**
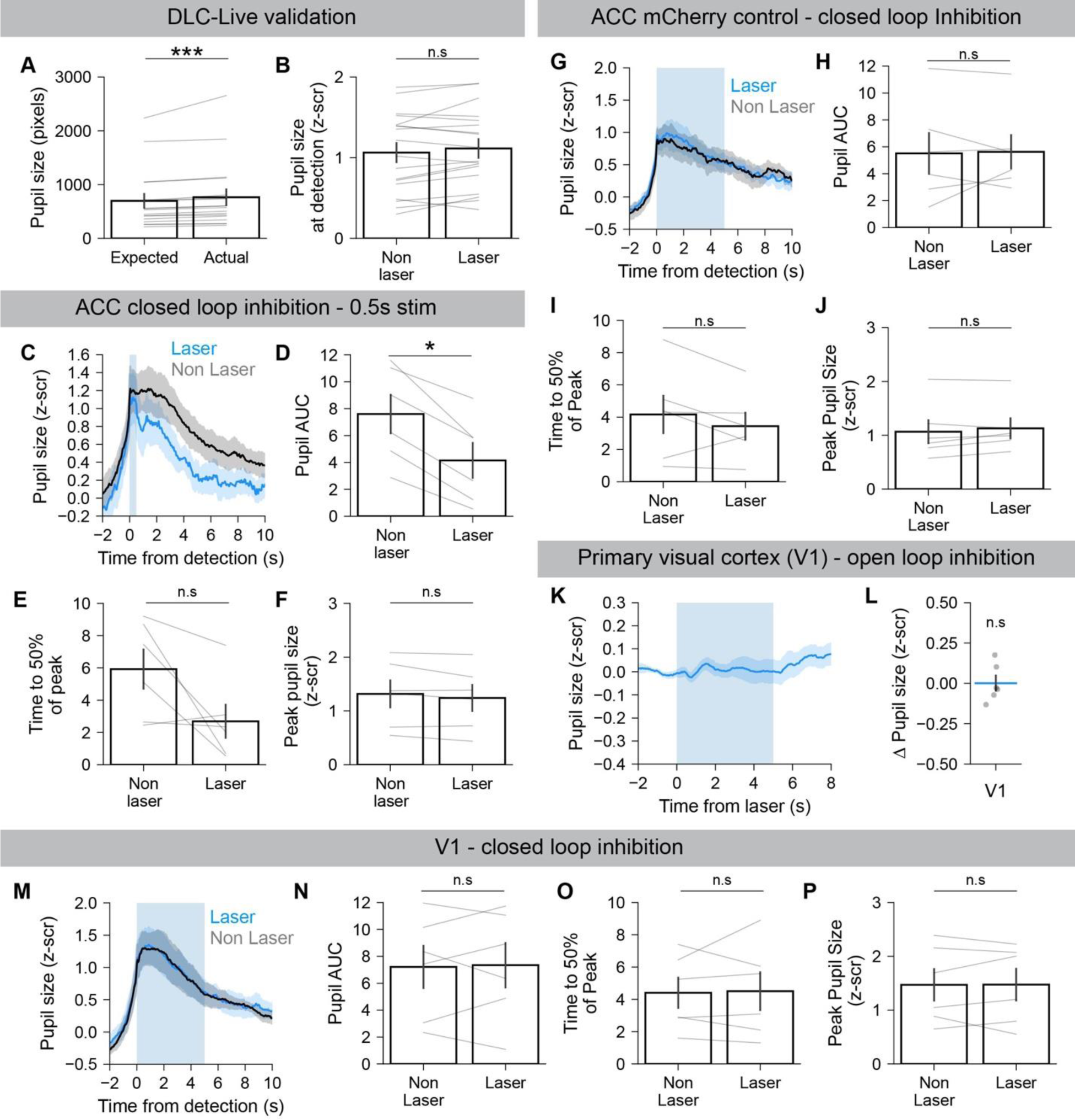
Closed loop optogenetic inactivation of the ACC, but not V1, suppresses ongoing pupil dilations. **(A)** Threshold value used for sessions (expected) compared to the average pupil size when dilation was detected (actual) (n = 18 mice; p < 0.0001, T = 0; Wilcoxon signed-rank test). **(B)** Average pupil size when dilation was detected for non-laser and laser trials (n = 18 mice; p = 0.14, T = 51; Wilcoxon signed-rank test). **(C)** Pupil size aligned to time of dilation detection for non-laser and laser trials. Blue shading shows photostimulation time for laser trials. For these experiments, a 0.5s laser duration was used. **(D)** Area under the curve (AUC) for non-laser and laser trials (n = 6 mice, p = 0.03, T = 0; Wilcoxon signed-rank test). **(E)** Time taken for pupil to decline to 50% of peak value (n = 6 mice, p = 0.09, T = 0; Wilcoxon signed-rank test). **(F)** Peak pupil size reached for laser and non-laser trials (n = 6 mice, p = 0.16, T = 0; Wilcoxon signed-rank test). **(G)** Same as C but for mCherry controls. For these experiments, a 5s laser duration was used. **(H)** Same as D but for mCherry controls (n = 6 mice, p = 1.0, T = 10; Wilcoxon signed-rank test). **(I)** Same as E but for mCherry controls (n = 6 mice, p = 0.16, T = 3; Wilcoxon signed-rank test). **(J)** Same as F but for mCherry controls (n = 6 mice, p = 0.22, T = 4; Wilcoxon signed-rank test). **(K)** Pupil size aligned to photostimulation onset for mice expressing VGAT-ChR2 in the primary visual cortex (V1; n = 6 mice). **(L)** Photostimulation evoked change in pupil size for mice shown in K (n = 6 mice, p = 1, T = 10; Wilcoxon signed-rank test against zero). **(M)** Same as C but for mice expressing VGAT-ChR2 in V1. For these experiments, a 5s laser duration was used. **(N)** Same as D but with V1 inactivation (n = 6 mice, p = 0.84, T = 9; Wilcoxon signed-rank test). **(O)** Same as E but for V1 inactivation (n = 6 mice, p = 1, T = 10; Wilcoxon signed-rank test). **(P)** Same as F but for V1 inactivation (n = 6 mice, p = 1, T = 10; Wilcoxon signed-rank test). All error bars are standard error of the mean.

**Extended Data Fig. 3:**
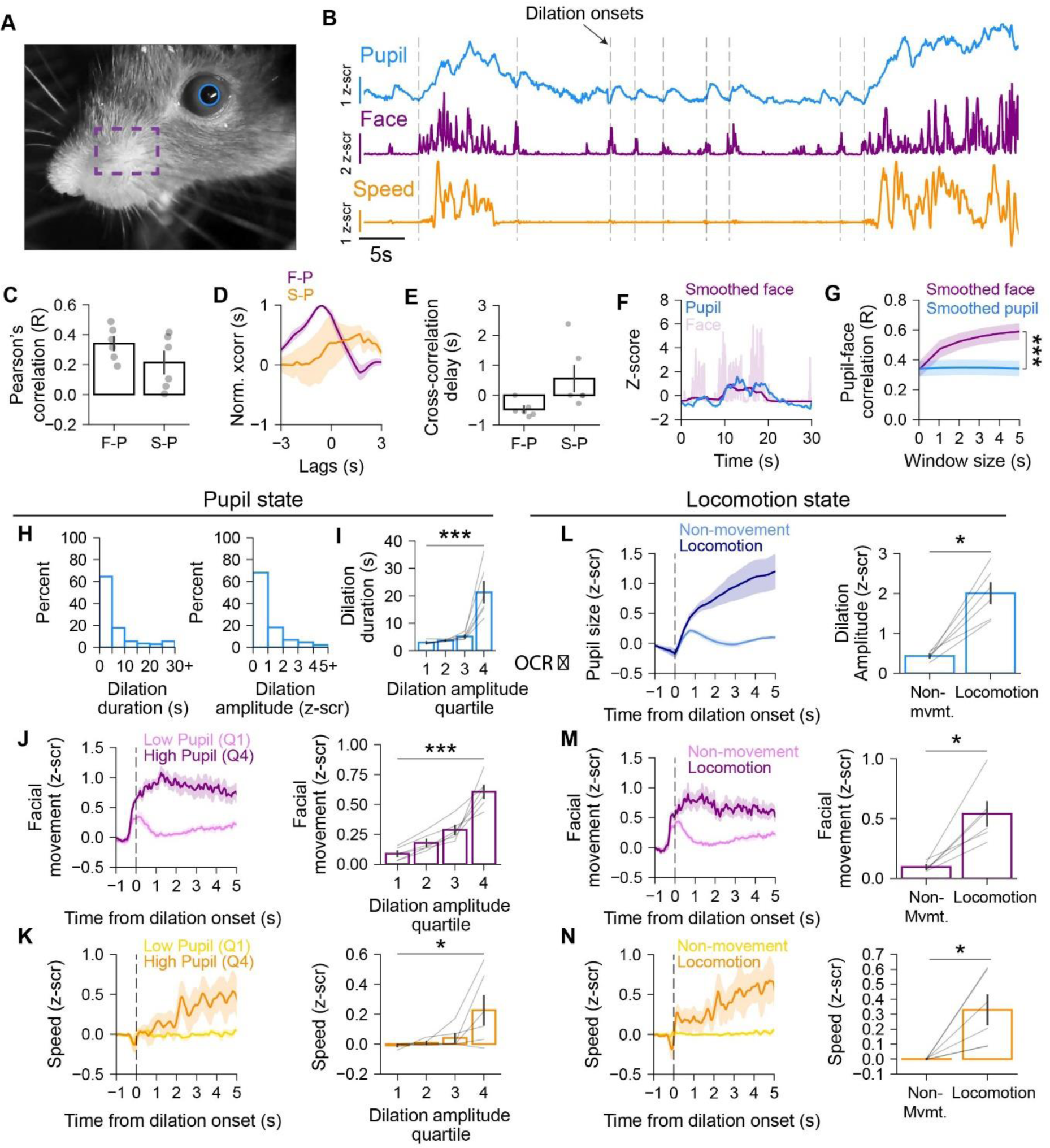
Analysis of behavioral metrics for arousal. **(A)** Example frame taken from a face video during a recording session. Blue outline shows the ellipse that was fit to the pupil using DeepLabCut tracked key points. Dashed purple rectangle shows the ROI used to quantify facial movement. **(B)** Example traces from a session showing pupil size, facial movement, and running speed. Dashed lines indicate onsets of pupil dilations. **(C)** Pearson’s correlation of facial movement (F) or running speed (S) with pupil size (P) (n = 6 mice). **(D, E)** Cross-correlation of changes in facial movement/running speed relative to pupil size. **(F)** Example trace showing traces of pupil size, and raw and smoothed face movements **(G)** Pearson’s correlation between facial movement and pupil size using different window sizes for smoothing either facial movement or pupil size. (n = 6 mice; main effect of window size F(5,25) = [77.83], p = 2e-14; main effect of smoothed condition F(1,5) = [125.43], p = 9.9e-5; interaction F(5,25) = [85], p = 7e-15; two-way repeated measures ANOVA). **(H)** *Left*, Histograms of pupil event durations (left) and amplitudes (right). **(I)** Pupil event durations across quartiles of pupil event amplitude (n = 6 mice; p = 0.0003, H = 19.04; Kruskal-Wallis test). **(J)** *Left*, facial movement aligned to onset of pupil dilation for pupil events in quartiles 1 (Q1) and 4 (Q4). *Right*, mean facial movement across quartiles of pupil event amplitude (n = 6 mice; p = 0.0003, H = 19.11; Kruskal-Wallis test). **(K)** *Left*, Running speed aligned to onset of Q1 and Q4 pupil dilation. *Right*, Mean running speed across quartiles of pupil event amplitudes (n = 6 mice; p = 0.027, H = 9.18; Kruskal-Wallis test). **(L)** *Left*, Pupil size aligned to onset of pupil dilations during non-movement and locomotion. *Right*, Pupil dilation amplitude for non-movement and locomotion conditions (n = 6 mice; p = 0.03, T = 0; Wilcoxon signed-rank test). **(M)** *Left*, traces of facial movement aligned to onset of non-movement and locomotion dilation events. *Right*, mean facial movement for non-movement and locomotion pupil events. (n = 6 mice; p = 0.03, T = 0; Wilcoxon signed-rank test). **(N)** *Left*, Running speed aligned to onset of pupil dilation for non-movement and locomotion pupil events. *Right*, mean wheel speed for non-movement and locomotion pupil events. (n = 6 mice; p = 0.03, T = 0; Wilcoxon signed-rank test). All error bars are standard error of the mean.

**Extended Data Fig. 4:**
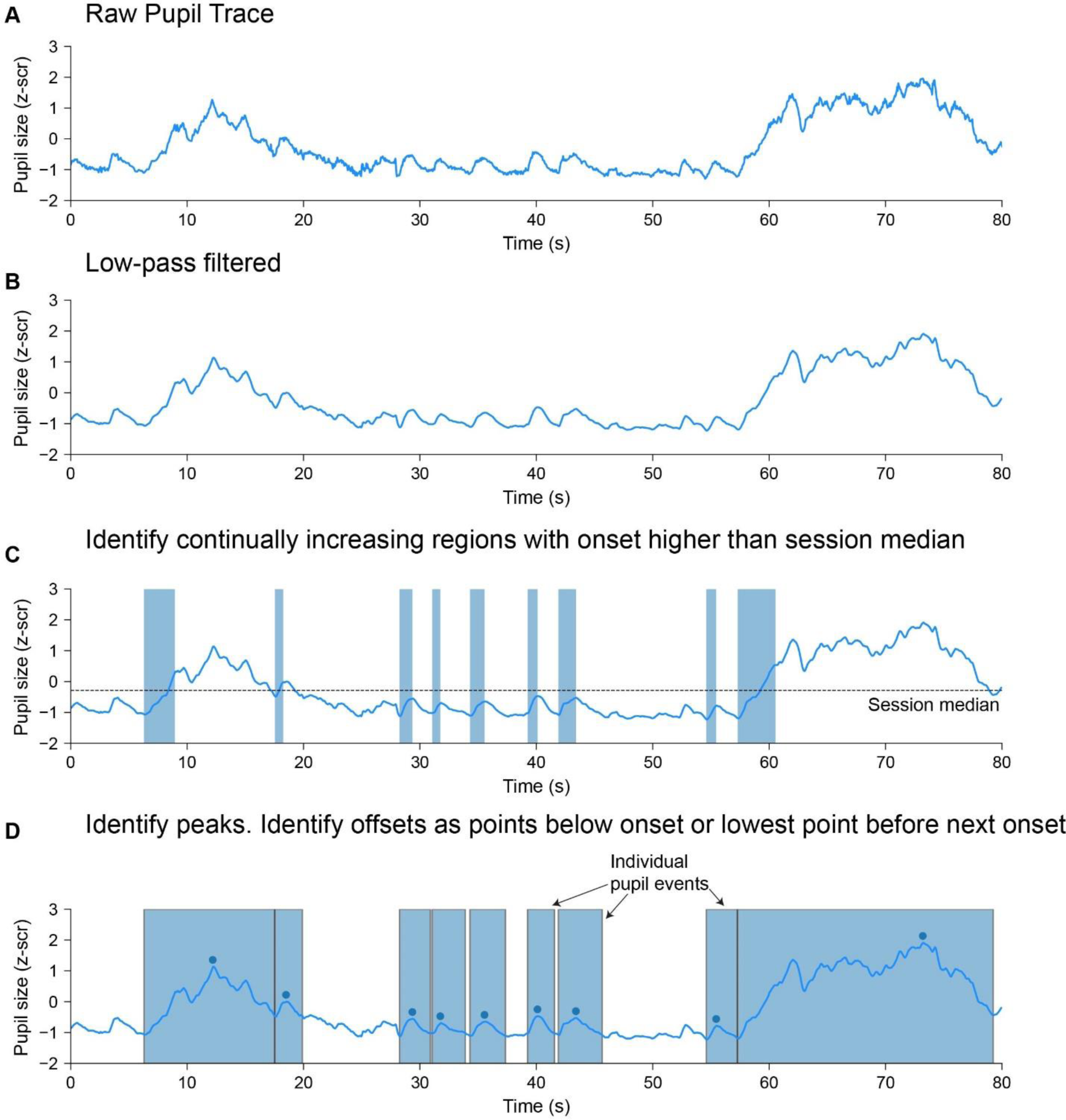
Method for detection of individual pupil dilation events. **(A)** Raw pupil trace taken from an example session. **(B)** Pupil trace after filtering with a 1st order low-pass Butterworth filter with a cutoff frequency of 1Hz. **(C)** Shaded blue areas show regions in which pupil size continually increases for at least 750ms. Dashed line shows the median pupil size for this session. Only regions with onset values below the session’s median pupil size were used. Regions within 1s of each other were merged. **(D)** The onset of the dilation event was taken to be the onset of the region. The offset of each event was taken to be the first point between the end of the region and the start of the next region where pupil size was lower than the onset pupil size. If pupil size did not decrease below the onset size in that window, offset was taken to be the lowest value in that window. Blue rectangles show individual pupil events. Blue circles show the maximum value or peak during each event.

**Extended Data Fig. 5:**
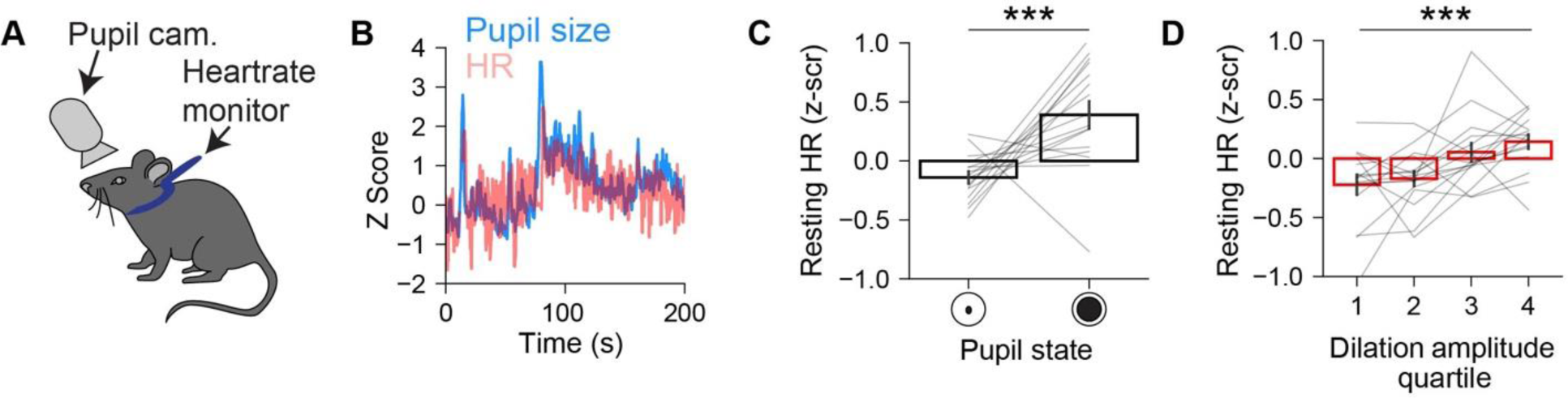
Pupil size is correlated with resting heart rate. **(A)** Schematic showing the experimental setup used to measure heart rate (HR) and pupil size. **(B)** Example trace of simultaneously recorded pupil size and HR. **(C)** Comparison of average HR when pupil is either dilated (z-score > 1.5) or constricted (z-score < 0). HR is significantly higher when pupil is dilated (n = 16 mice, p = 0.004, T = 15; Wilcoxon signed-rank test). **(D)** Average HR across quartiles of pupil dilation amplitude (n = 16 mice; p = 0.0009, H = 16.46; Kruskal-Wallis test). All error bars are standard error of the mean.

**Extended Data Fig. 6:**
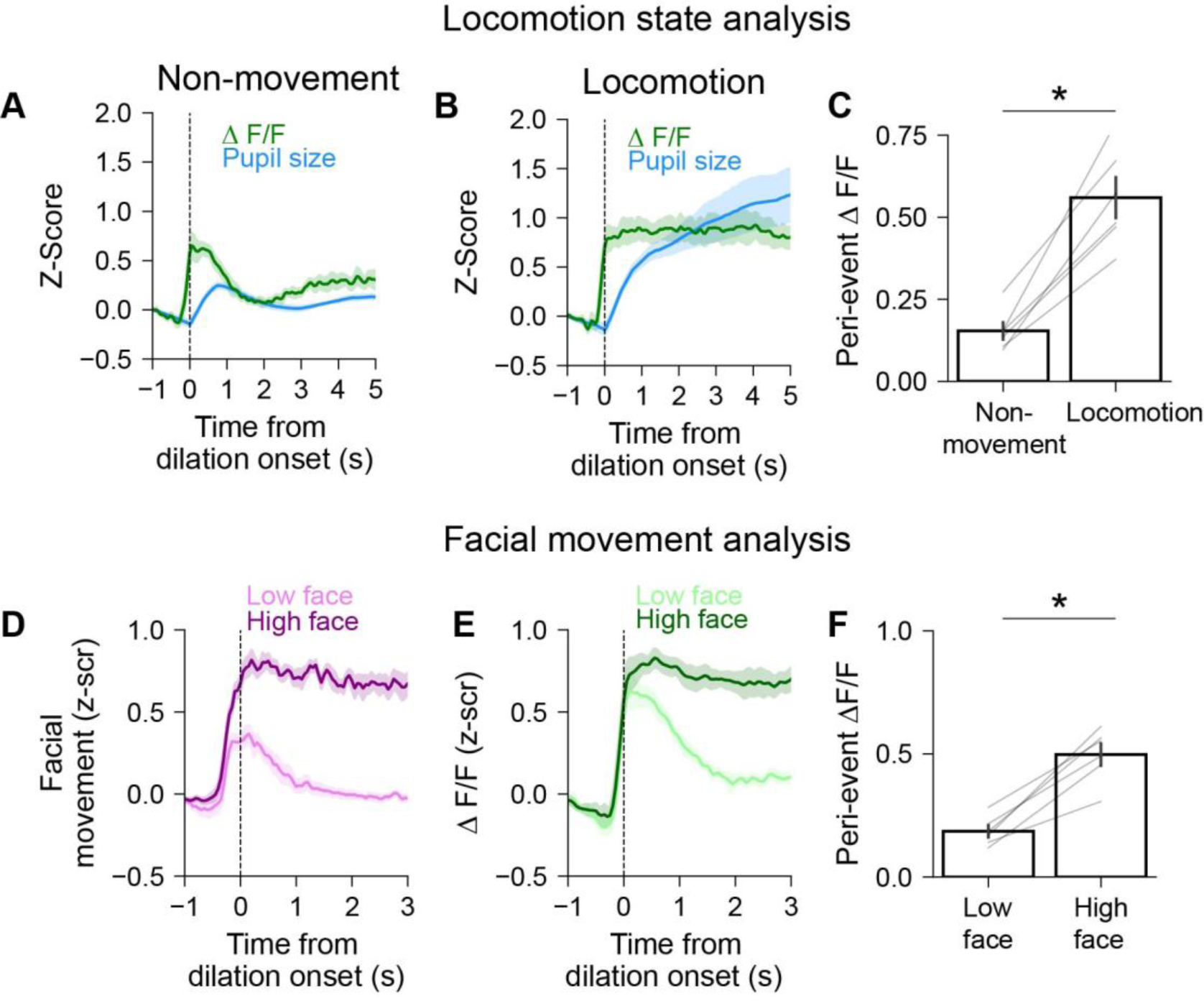
Relationship of ACC activity to locomotion and facial movements. **(A)** For all non-movement dilation events, pupil size (blue) and ACC activity (green) aligned to onset of pupil dilation (n = 6 mice). **(B)** Same as A but for locomotion dilation events (n = 6 mice). **(C)** Pupil related ACC activity during non-movement and locomotion (n = 6 mice; p = 0.03, T = 0; Wilcoxon signed-rank test). **(D)** Facial movement aligned to onset of pupil dilation for events with low or high facial movement (determined by median split) (n= 6 mice). **(E)** Same as D but for ACC activity (n = 6 mice). **(F)** ACC activity during events with low or high facial movement (n = 6 mice; p = 0.03, T = 0; Wilcoxon signed-rank test).

**Extended Data Fig. 7:**
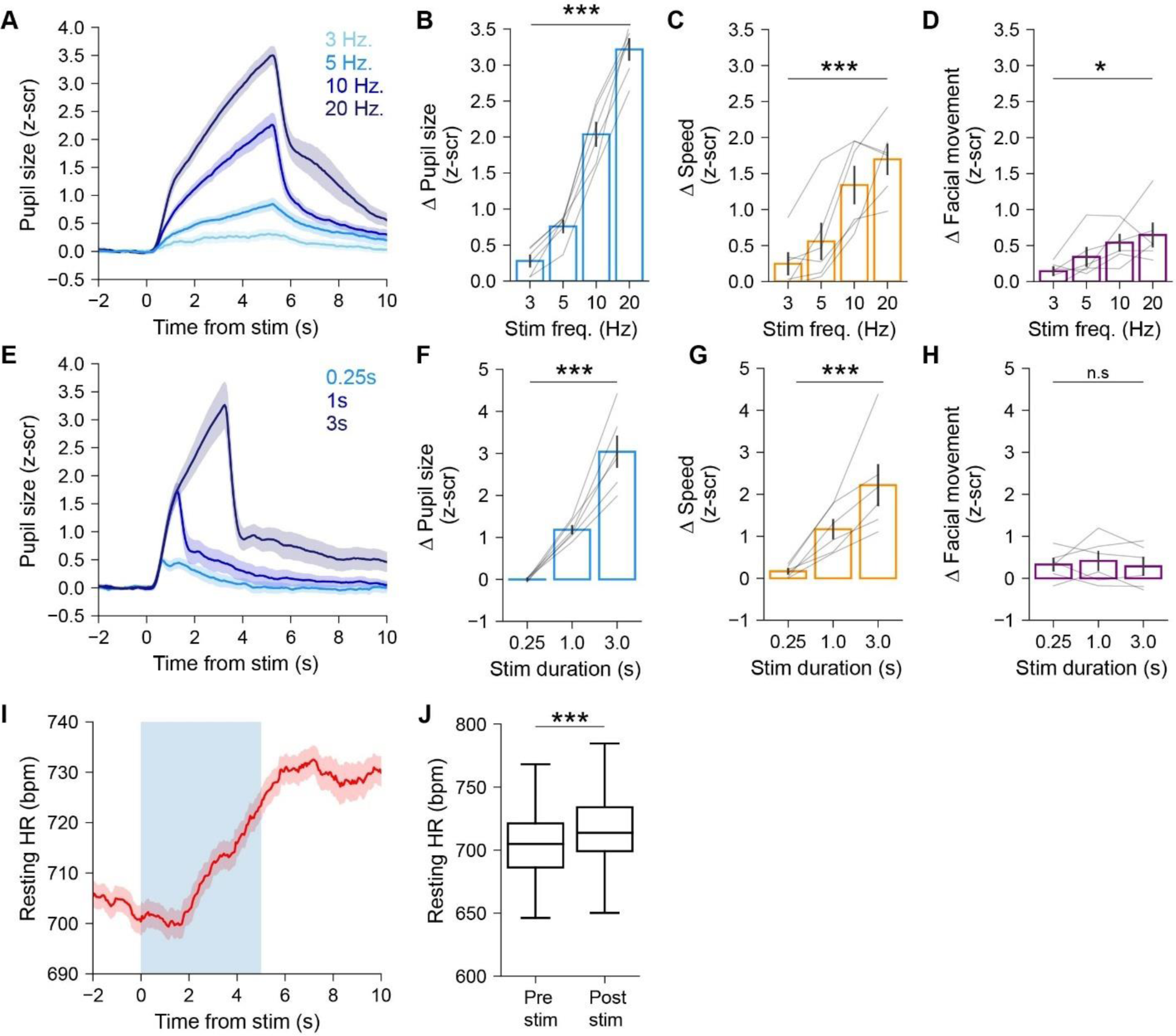
Effect of optogenetic ACC on pupil size, facial movements, locomotion, and heart rate. **(A)** Pupil size aligned to photostimulation at variable frequencies stimulation frequencies. **(B)** Change in pupil size with varying photostimulation frequencies (n = 6 mice; p = 0.0001, H = 20.94; Kruskal-Wallis test). **(C)** Same as B but for locomotion speed (n = 6 mice; p = 0.003, H = 13.99; Kruskal-Wallis test). **(D)** Same as B but for facial movement (n = 6 mice; p = 0.016, H = 10.35; Kruskal-Wallis test). **(E)** Pupil size aligned to photostimulation onset for varying durations. **(F)** Change in pupil size with varying photostimulation durations (n = 6 mice; p = 0.0005, H = 15.16; Kruskal-Wallis test). **(G)** Same as F but for locomotion speed (n = 6 mice; p = 0.001, H = 13.05; Kruskal-Wallis test). **(H)** Same as F but for facial movement (n = 6 mice; p = 0.83, H = 0.36; Kruskal-Wallis test). **(I)** Heartrate aligned to photostimulation onset (n = 171 trials from 3 mice). **(J)** Pre-stim heartrate versus post-stim heartrate (n = 171 trials from 3 mice, p <0.00001, T = 2587; Wilcoxon signed-rank test). All error bars are standard error of the mean.

**Extended Data Fig. 8:**
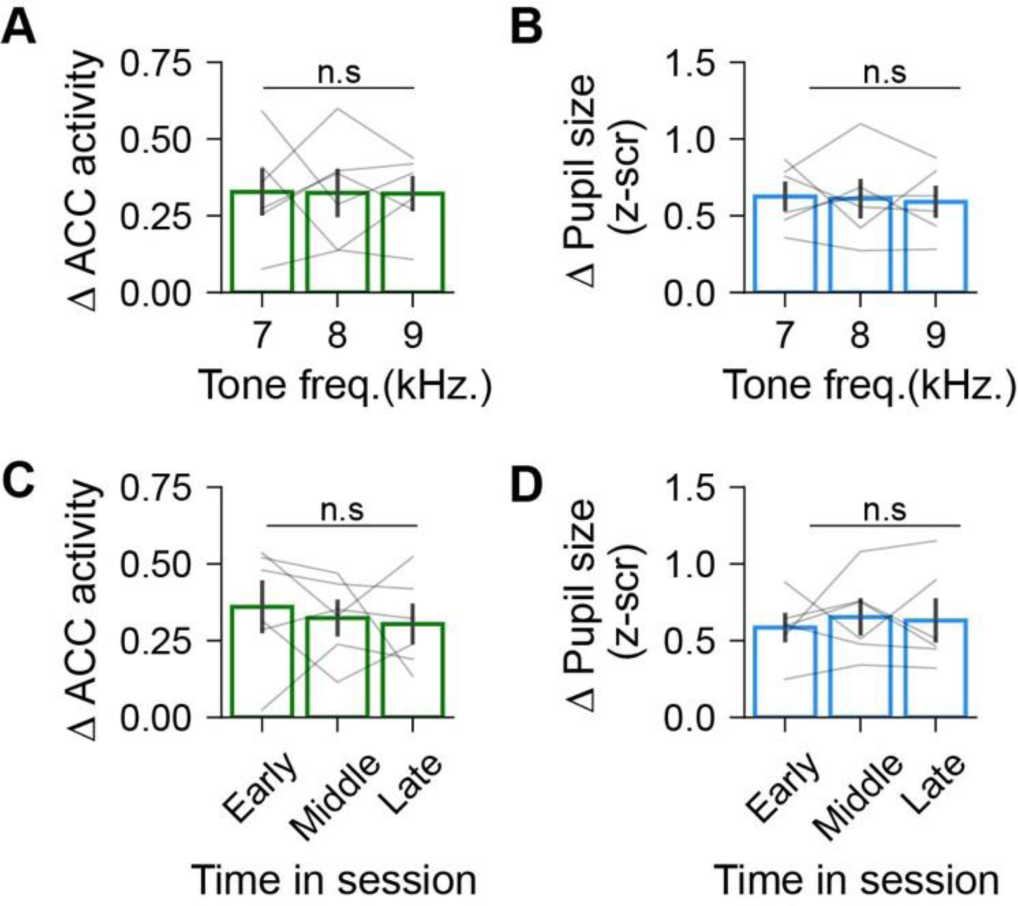
ACC activity and pupil size do not depend on tone frequency or habituation to auditory stimuli. **(A)** Tone-evoked ACC activity for each auditory tone frequency. Evoked ACC activity responses were not dependent on tone frequency (n = 6 mice; p = 0.95, H = 0.11; Kruskal-Wallis test). **(B)** Same as A but for tone-evoked changes in pupil size (n = 6 mice; p = 0.98, H = 0.05; Kruskal-Wallis test). **(C)** Tone-evoked changes in ACC activity at different points throughout session. ACC activity did not show habituation to the auditory tones (n = 6 mice; p = 0.74, H = 0.61; Kruskal-Wallis test). **(D)** Tone-evoked changes in pupil size at different points throughout session. Pupil size did not show habituation to the auditory tones (n = 6 mice; p = 0.93, H = 0.15; Kruskal-Wallis test). All error bars are standard error of the mean.

**Extended Data Fig. 9:**
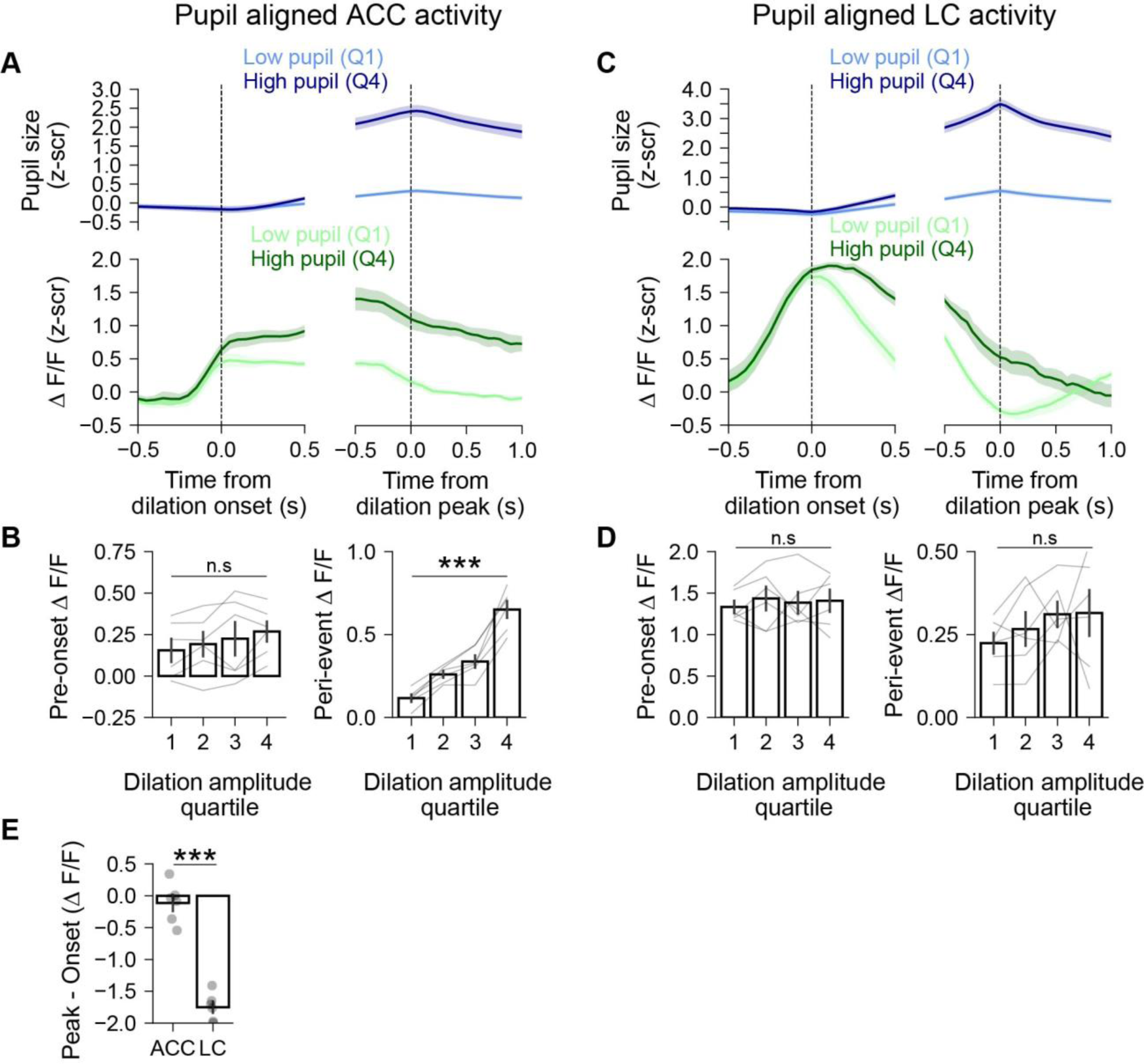
ACC and LC activity differentially scales with pupil size. **(A)** *Top*, Pupil size aligned to onset (left) and peak (right) of pupil dilations for events in quartiles 1 (Q1) and 4 (Q4). *Bottom*, same as top but showing ACC activity(n = 6 mice). **(B)** *Left*, ACC activity before pupil dilation onset shown across quartiles of pupil event amplitude (n = 6 mice; p = 0.69, H = 1.49; Kruskal-Wallis test). *Right*, ACC activity during dilation events across quartiles of pupil event amplitudes (n = 6 mice; p = 0.0002, H = 20.25; Kruskal-Wallis test). **(C)** Same as A but for LC recordings (n = 4 mice). **(D)** *Left*, Pre-onset LC activity across quartiles of pupil event amplitudes (n = 6 mice; p = 0.96, H = 0.29; Kruskal-Wallis test). *Right*, LC activity during dilation events across quartiles of pupil event amplitude (n = 6 mice; p = 0.60 H = 1.85; Kruskal-Wallis test). **(E)** Difference between activity at peak of dilation versus onset of dilation in ACC or LC (n = 6 LC mice, 6 ACC mice, p = 0.005, U = 0; Mann-Whitney U rank test).

## Acknowledgements

This work was supported by grants from the National Institute of Mental Health (R00-MH112855 to R.H.), National Institute of Alcohol Abuse and Alcoholism and the NIH Office of the Director (R01-AA030594), Brain and Behavior Research Foundation (NARSAD Young Investigator Award to R.H. and V.B.-P.), Brain Canada (Future Leaders in Canadian Brain Research to V.B.-P.), the Natural Sciences and Engineering Research Council of Canada (Discovery Grant RGPIN-2021-03284 to V.B.-P.), a New Frontiers in Research Fund (NFRFE-2022-00342 to V.B.-P.), Fonds de recherche du Québec, Santé (Research Scholars - Junior 1 Salary Award to V.B.-P.). We thank Drs. Cherish Ardinger and Wes Evans and other members of the laboratory for providing critical feedback.

## Author contributions

N. Chintalacheruvu and R. Huda designed experiments, A. Kalelkar performed surgeries and histological experiments, N. Chintalacheruvu performed and analyzed ACC optogenetic and fiber photometry experiments, J. Boutin performed LC fiber photometry experiments, and N. Chintalacheruvu and V. Breton-Provencher analyzed LC fiber photometry experiments. N. Chintalacheruvu and R. Huda wrote the manuscript with input from all authors. R. Huda supervised the study. R. Huda and V. Breton-Provencher secured funding.

## Methods

### Animals

Behavioral experiments were performed on male and female C57/BL6J mice maintained on a reversed light/dark circadian cycle with ad libitum access to standard mouse chow and water. Mice of either sex were at least 10 weeks old at the start of behavioral experiments. Optogenetic activation of ACC GABAergic neurons were done using transgenic VGAT-Cre mice. All animal procedures were performed in strict accordance with protocols approved by the Rutgers Comparative Medicine Resources and conformed to NIH standards.

### Surgical procedures

Surgical methods were as described previously^41,42^. Briefly, all animals received stereotactic viral injection, optic fiber implant, and headplate implant. Surgeries were performed under isoflurane anesthesia (4% induction, 1-3% maintenance). Body temperature was maintained at 37.5◦C via a heating pad integrated into the base of the stereotaxic frame and a temperature controller (53800, Stoelting). Mice were given a subcutaneous injection of extended-release buprenorphine (Ethiqa-XR, 3.25mg/kg) before surgery to provide analgesia for up to 72 hours post-surgery; meloxicam (10 mg/kg) was provided if additional analgesia was required during the recovery period. Anesthetized mice were head-fixed in a stereotaxic frame (51500D, 140 Stoelting). Scalp hair was removed using a depilatory cream (Nair) and the scalp was disinfected using 3x alternating scrubs with betadine and 70% ethanol solution. A portion of the scalp was removed, and conjunctive tissues cleared after treatment with 3% hydrogen peroxide. The skull was abraded with a dental drill to improve adhesion of dental cement.

For viral injections, a burr hole was made in the skull above the injection site (ACC: 0.3mm AP, 0.3mm ML, −0.9mm DV; V1: −3.5mm AP, 2.5mm ML, −0.5mm DV; LC: −5.1mm AP, 0.9mm ML, −2.8mm DV). Viruses were pressure injected using a Nanoject III (Drummond) or a Nanoliter 2020 Injector (WPI) through a glass pipette slowly lowered into the craniotomy. The following viruses were used at the specified injection volumes: AAV1-syn-jGCaMP8m-WPRE, 120nL (Addgene catalog # 162375); AAV9-syn-FLEX-jGCaMP8s-WPRE (Addgene catalog #162377-AAV9), 300 nL; AAV5-EF1a-double floxed-hChR2(H134R)-mCherry-WPRE-HGHpA, 120nL (Addgene catalog # 20297); AAV5-CaMKIIa-hChR2(H134R)-mCherry, 100-120nL (Addgene catalog # 26975); and AAV5-CaMKIIa-mCherry (Addgene catalog # 114469), 120nL.

Following virus injections, a fiber optic was slowly inserted into the brain ∼0.1mm above the injection site. Unilateral fiber optic cannulae were used for fiber photometry experiments (400µm core, 0.5 NA, RWD Life Science) and custom-made bilateral cannulae were implanted for ACC optogenetic experiments (200µm, 0.39NA, 0.7mm distance between fiber optics, Doric Lenses). For optogenetic experiments in V1, single fibers were implanted in each hemisphere (200µm, 0.39NA, RWD). The fiber optic cannula and a headplate was secured adhered to the skull using Metabond dental cement (C&B)

Animals were allowed to recover in their home cage over a warm water blanket until they recovered from anesthesia. Moistened food chow and hydrogel was provided. Animals were monitored post-operatively for 3-4 days. Mice were singly housed for the remainder of the experiment and recovered from surgery for 3-8 weeks before beginning experiments.

### Quantification of behavioral variables

During all behavioral experiments, an infrared camera was used to record videos of the mouse’s face including the pupil. An infrared light source (LN-3, ORDRO) was used to illuminate the face, and an ambient light source was used to keep the pupil sufficiently constricted (lux). Closed loop optogenetic experiments were done using a USB Camera (Day & Night Vision, ArduCam) and separate LED light (Book Light, Vont). All other experiments were done using an OpenMV camera (H7) with ambient light coming from an LED on the camera. Pupil videos were then processed using DeepLabCut to track eight points on the perimeter of the pupil, and frames with low confidence estimations (<0.95) were dropped. Least squares fitting was used to fit the eight points to an ellipse. Pupil size was then quantified by finding the area of this ellipse. Finally, pupil measurements were interpolated at 20hz. To quantify facial movement, an ROI was drawn over the whisker pad, and facial movements were quantified as the difference in average pixel intensity between ROIs of consecutive frames. Facial movements were interpolated at 20hz. In experiments that included locomotion measurements, animals were head fixed onto a wheel attached to a rotary encoder (Yumo). Locomotion speed was sampled at a rate of 20 Hz. Animals were habituated to the entire head-fixed setup including the wheel for at least 3 days before their first session.

### Optogenetic stimulation

Bilateral photostimulation of CaMKII-ChR2 expressing ACC neurons was performed by connecting the output of two 470nm fiber coupled LEDs (M470F3, Thorlabs) to a dual fiber optic patch cord (Doric Lenses). The patch cord was connected to the mouse’s ferrule implant using a sleeve. Bilateral optogenetic stimulation in VGAT-cre mice was done using a 470nm laser (MBL-III-470, Optoengine). The laser was coupled to a beam splitter (Doric Lenses mini cube), the output of which was connected to the dual fiber patch cord. The patch cord was then connected to the implanted ferrule with a coupling sleeve (Doric Lenses or RWD). For these experiments, black electrical tape was wrapped around the sleeve to prevent light leakage. For stimulation of CaMKII-ChR2 expressing ACC neurons, combined light power at the end of the fiber tip was ∼1 mw. Higher light intensity (10 - 12 mw) was used for stimulation of ACC GABAergic neurons.

Open-loop optogenetic experiments were carried out using custom-made MATLAB and Arduino scripts. The MATLAB and Arduino interface was used to synchronize timings of behavioral variables and optogenetic stimulation. Stimulation frequency and duration were specified by the MATLAB script. For optogenetic experiments involving stimulation of CaMKII ACC neurons, sessions were carried out using 10ms pulses either at different durations (250ms, 1000ms, 3000ms) or different frequencies (3hz, 5hz, 10hz, 20hz). Sessions using different durations were done using a 20hz stimulation, and sessions using different frequencies were done using 5s stimulation. All sessions were 30 minutes long and had an inter-trial-interval of 20s. For experiments in Fig. 4, additional sessions were carried out using a single 10ms pulse with an inter-trial-interval of 10s.

For optogenetic experiments involving stimulation of ACC GABAergic neurons, sessions were carried out using 10ms pulses at 20Hz with a 5s train duration. Additional sessions were carried out either at different durations (500ms, 1000ms, 2000ms) or different frequencies (5hz, 10hz, 20hz, 30hz). Sessions using different durations were done using a 20hz stimulation, and sessions using different frequencies were done using 5s stimulation. All sessions were 30 minutes long and had an inter-trial-interval of 20s.

Closed loop optogenetic experiments were performed using DeepLabCut-Live! GUI^36^, a software package for real-time pose estimation. DeepLabCut model trained for offline pupil size quantification was used for online detection. A custom-made Python script integrated with DeepLabCut-Live! GUI was used to trigger optogenetic stimulation based on detection of real-time pupil dilation events. A pupil dilation event was detected each time pupil size increased beyond a preset threshold pupil size. On 50% of detected trials, optogenetic stimulation was given. The threshold pupil size was determined through a calibration session (10-20 min) in which the mouse’s pupil was recorded. Pupil traces were z-scored and peaks in pupil size were then found using the Python function *findpeaks* with a prominence of 2 z-scores. The threshold was set to be 25% of the average prominence value of detected peaks. The threshold z-scored value was converted back to pixel size and used as the threshold for identifying pupil dilations for each session. For analysis of closed loop experiments, only trials where the slope of the pupil size preceding trial onset was positive were used.

### Fiber photometry

Fiber photometry recordings of bulk ACC calcium activity were carried out using the Tucker-Davis Technologies RZ10X system and Synapse software. Excitation light was sinusoidally modulated for 465nm (330 Hz) and 405nm (210 Hz) wavelengths used for obtaining calcium-dependent and isosbestic emission signals, respectively. Excitation light and emitted light from the sample was routed to/from the TDT system and the animal with a fluorescence minicube (Doric Lenses). Videos of the face were taken with an OpenMV camera (H7) running custom micropython scripts at 20 fps. Simultaneously, an Arduino Uno was used to sample wheel speeds from a rotary encoder at 20hz. A custom MATLAB script was used to record the wheel speeds from the Arduino. To ensure synchronization of fiber photometry recordings and behavioral measurements, each time a video frame was acquired, a TTL signal was sent to the Synapse software. Each time the Arduino sampled running speed, a separate TTL signal was sent to the Synapse software. The timepoints for each stream of TTL signals were used to interpolate each behavioral signal at 20Hz. This ensured that facial videos, wheel speeds, and photometry signals were synchronized to the same clock. For analysis, ΔF/F (see below) was resampled to 20hz at the interpolated time points for wheel speed and facial videos.

For LC recordings, we used a custom-built, camera based system adapted from a previously published design^43^. Excitation light from 405 and 470 nm wavelengths were coupled into the microscope and interleaved at a rate of 40 Hz, for a final acquisition rate of 20 Hz. For both recordings, a ceramic coupling sleeve (Doric Lenses) was used to couple the patch cord to the implant. Videos of the face were taken with a Blackfly Camera (BFS-U3-04S2M-CS) using the Spinview software. To synchronize face videos with calcium recordings, a custom Python script was used to send a TTL signal to the photometry system. The same TTL signal also toggled a red LED placed in the camera field of view. Video and calcium recordings were synchronized by aligning the onset of the TTL signal with the onset of the red light in the face recording.

A custom Python script was used to process photometry recordings. For calculating ΔF/F, a first degree least squares polynomial was used to fit the isosbestic channel (405nm excitation) to the signal channel (465nm excitation). ΔF/F was calculated as 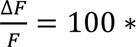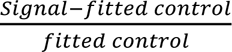

### Auditory tones

Auditory stimuli were created using the MATLAB package PsychToolBox-3 and presented using dual speakers (Pebble V2, Creative) placed approximately 20 cm from the mouse. For all experiments, sound intensity was measured to be 80dB using an SPL meter (CM-130, Galaxy Audio). Tone frequencies randomly varied throughout the session and tone duration was set at 0.5s. For optogenetic experiments, frequencies of 8khz, 9khz and 10khz were used with an interstimulus interval of 20 seconds. For fiber photometry experiments, frequencies of 7khz, 8khz and 9khz were used with a random interstimulus interval drawn from an exponential distribution with a mean of 15s with cutoffs of 10s and 20s. For optogenetic experiments, laser stimulation started 5 seconds before tone onset and lasted for 5 seconds after tone onset (10 seconds total laser duration). For fiber photometry recordings, analysis was restricted to trials where there was an increase in pupil size following auditory tones relative to the preceding baseline.

### Heartrate measurement

Heartrate was measured using a pulse oximeter (Starr Life Sciences). Mice were briefly anesthetized with isoflurane (4% induction, 1-3% maintenance). Depilatory cream (Nair) was used to remove hair from the mouse’s neck. Experimental sessions were performed at least 1 day following anesthesia. Head-fixed mice were fitted with a collar sensor to collect oximetry signals from the carotid artery. During recording, heartrate samples were sent to an Arduino as an analog input using an Analog Data Output Module (Starr Life Sciences). To synchronize measurements and adjust for lag between heartrate measurements and pupil recordings, heartrate measurements were shifted backwards by 1.57s. To convert analog inputs to heartrate, inputs were mapped to 0V to 5V and multiplied by a scalar (200). The Analog Data Output Module also sent inputs to the Arduino regarding the error status of each heartrate sample. Samples with pulse related error codes were dropped.

### Pupil event classification

Detection of pupil dilation events was adapted from a recent method ^44^ and done through the following steps: 1) Pupil trace was filtered using a 1st order low-pass Butterworth filter with a cutoff frequency of 1hz; 2) Regions in which pupil size was continually increasing for at least 750ms were identified. Only regions whose onset value was below the session’s median pupil size were used. Regions within 1s of each other were considered to be one region; 3) The onset of the dilation event was taken to be the onset of the region. The offset of each event was taken to be the first point between the end of the region and the start of the next region where pupil size was lower than the onset pupil size. If pupil size did not decrease below the onset size in that window, offset was taken to be the lowest value in that window; 4) Event duration was taken to be the time between onset and offset. Event amplitude was taken to be the maximum pupil size between onset and offset subtracted by the onset pupil size.

### Classification of locomotion and pupil state

For a subset of analyses, pupil dilations were classified as non-movement or running dilation events. Since the max running speed varied between mice, the speed thresholds used to classify events were calculated for each session. Running dilation events included pupil dilations for which the mean running speed during the dilation (defined by time of onset to offset) was greater than 65th percentile of running speed recorded in the session. Quiet dilation events included pupil dilations for which the absolute value of mean running speed during the dilation was less than the 45th percentile of running speed. The absolute value of running speed was used here to ensure that events with little movement in either the forward or backwards direction were identified.

In Fig. 4, trials were classified as non-locomotion trials or locomotion trials based on the mean running speed in a −0.5s to 2s window around each trial onset. Trials with a mean running speed less than 1AU were classified as non-locomotion trials, while trials with a mean running speed greater than 5AU were classified as locomotion trials.

In Extended Data Figs. 3 and 9, pupil dilation events were split into quartiles. Importantly, quartiles were computed within individual sessions. For clarity, plots show traces aligned to only the first and fourth quartile.

### Other analyses

Trial aligned z-scored traces were baseline corrected except in Figs. 1, 4A, 4D, 5I and Extended Data Fig. 2. In Extended Data Fig. 3, running speed, pupil size and facial movement were aligned to the onsets of pupil dilation. For each mouse, the session-averaged onset aligned trace was calculated (window: −2s to 2s) and used for normalized cross correlation analysis.

In Fig.6, we quantified the onset of neural activity in relation to the onset of pupil dilation in either the ACC or LC. To account for high levels of noise in single trials, onsets were calculated based on traces of session-averaged pupil sized aligned to the time of dilation onsets. Activity onset was defined as the first ΔF/F point in each session-averaged trace where activity increased continuously for at least 0.25s. The t_50_ was also quantified using the session-averaged trace. To calculate the t_50_, session-averaged traces were first baseline corrected by subtracting the minimum. The t_50_ was defined as the timepoint relative to pupil dilation onset where activity reached 50% of its maximum value.

